# Locomotor-related propriospinal V3 neurons produce primary afferent depolarization and modulate sensory transmission to motoneurons

**DOI:** 10.1101/2022.07.04.498712

**Authors:** Shihao Lin, Krishnapriya Hari, Ana M. Lucas-Osma, Sophie Black, Aysan Khatmi, Karim Fouad, Monica A. Gorassini, Yaqing Li, Keith K. Fenrich, David J. Bennett

## Abstract

When a muscle is stretched it not only responds with a reflex, but the sensory afferent feedback also depolarizes many afferents throughout the spinal cord (termed primary afferent depolarization, PAD), readying the whole limb for further disturbances. This sensory-evoked PAD is thought to be caused by a trisynaptic circuit, where sensory input activates first order excitatory neurons that activate GABAergic neurons that in turn activate GABA_A_ receptors on afferents to cause PAD, though the identity of these first order neurons is unclear. Here we show that these first order neurons are propriospinal V3 neurons, since they receive extensive sensory input and in turn innervate GABAergic neurons that cause PAD, because optogenetic activation or inhibition of V3 neurons in mice mimics or inhibits sensory-evoked PAD, respectively. Furthermore, persistent inward sodium currents (Na PICs) intrinsic to V3 neurons enable them to respond to transient inputs with long-lasting responses, explaining the long time-course of PAD. Also, local optogenetic activation of V3 neurons at one segment causes PAD in other segments, due to the long propriospinal tracts of these neurons, explaining the widespread radiation of PAD across the spinal cord. This in turn facilitates monosynaptic reflex transmission to motoneurons across the spinal cord. Additionally, we find that V3 neurons directly innervate proprioceptive afferents, causing a glutamate receptor mediated PAD (glutamate PAD). Finally, we show that increasing the spinal cord excitability with either GABA_A_ receptor blockers or chronic spinal cord injury causes an increase in the glutamate PAD, perhaps contributing to spasms after SCI. Overall, we show the V3 neuron has a prominent role in modulating sensory transmission, in addition to its previously described role in locomotion.

## INTRODUCTION

Sensory feedback is essential for maintaining posture and producing accurate movement (Capaday & Stein, 1986; Bennett *et al*., 1994; Bennett *et al*., 1996; Rossignol *et al*., 2006), without which incoordination, joint injury and falling occur, especially with peripheral neuropathies, aging and spinal cord injuries (Andrechek *et al*., 2002; Bouyer & Rossignol, 2003; Rossignol *et al*., 2006; Ahmad *et al*., 2019; Estilow *et al*., 2019; Takeoka & Arber, 2019). Accordingly, the spinal cord has evolved complex sensory control systems to integrate the many sensory inputs that come from the limbs into the motor systems (Rossignol *et al*., 2006; Gosgnach *et al*., 2017; Ziskind-Conhaim & Hochman, 2017; Ueno *et al*., 2018; Lalonde & Bui, 2020; Zholudeva *et al*., 2021), allowing appropriate reflexes to support movements and respond rapidly to external disturbances, like when tripping or falling. The first line of sensory control is a direct modulation of the sensory afferents themselves within the spinal cord, by axoaxonic connections onto afferents from specialized GABAergic neurons (GAD2_+_, abbreviated here GABA_axo_ neurons)(Hughes *et al*., 2005; Betley *et al*., 2009; Fink, 2013; Fink *et al*., 2014; Hari *et al*., 2021). This activates GABAA receptors on afferents, which ultimately leads to a large depolarization, termed primary afferent depolarization (PAD), mediated by outward chloride currents produced by the unusually high intracellular chloride concentrations in sensory neurons, compared to in other adult neurons (Barron & Matthews, 1938; Rudomin & Schmidt, 1999; Szabadics *et al*., 2006; Bardoni *et al*., 2013; Lucas-Osma *et al*., 2018).

While the existence of PAD has been known for nearly a century (Barron & Matthews, 1938), the underlying neuronal circuits that cause it are uncertain and its function has recently been disputed, especially with the advent optogenetic methods that allow direct activation of the neurons causing PAD (Lucas-Osma *et al*., 2018; Hari *et al*., 2021). For example, while optogenetic activation of GABA_axo_ neurons causes GABAA-mediated PAD, this PAD does not account for presynaptic inhibition of proprioceptive sensory feedback to motoneurons (Hari *et al*., 2021). Instead, activation of GABAB receptors on afferent terminals causes presynaptic inhibition independently of PAD, since it is blocked by GABAB antagonists, and mainly only GABAB, and not GABAA, receptors are expressed at proprioceptive afferent terminals near motoneurons (Curtis & Lacey, 1994; Fink, 2013; Hari *et al*., 2021). Furthermore, PAD in proprioceptive afferents has been shown to be caused by GABA_axo_ neuron contacts onto GABAA receptors at or near nodes of Ranvier in the many myelinated branches of these afferents within the spinal cord, completely contrary to the long-standing assumption that PAD arises from afferent terminals and produces presynaptic inhibition (Eccles *et al*., 1961; Eccles *et al*., 1962; Lucas-Osma *et al*., 2018; Hari *et al*., 2021). This nodal GABA action paradoxically facilitates spike propagation failure at branch points (termed nodal facilitation), which is otherwise common without GABA innervation (Hari *et al*., 2021). How this complex mixture of inhibition and facilitation is regulated by GABA_axo_ neurons is even more uncertain, and it may well differ at rest compared to during locomotion (Gossard, 1996). Further, we know little about even the neurons that directly innervate GABA_axo_ neurons that cause PAD in group Ia proprioceptive afferents, though considerable progress has recently been made in understanding the similar circuits that mediate PAD in low threshold mechanoreceptors (e.g. CCK+ neurons innervate GABAergic neurons)(Lidierth & Wall, 1998; Koch *et al*., 2017; Zimmerman *et al*., 2019).

PAD and associated GABA_axo_ neuron activity, are well established to be tightly regulated during tasks like resting postural maintenance, walking and reaching, but how and why this occurs remain unclear (Gossard, 1996; Rudomin & Schmidt, 1999; Rossignol *et al*., 2006; Ueno *et al*., 2018; Moreno-Lopez *et al*., 2021). We know that when a muscle at rest is stretched it not only responds with a reflex, but the proprioceptive and cutaneous feedback caused by the imposed movement elicits a PAD in many afferents, readying the whole body for further disturbances, though the balance of presynaptic inhibition and nodal facilitation remains unclear (Rudomin & Schmidt, 1999; Hari *et al*., 2021). The PAD evoked by such sensory feedback is thought to be caused by a minimally trisynaptic circuit, where sensory inputs activate first order excitatory neurons that activates GABA_axo_ neurons that in turn produce PAD in afferents (Rudomin & Schmidt, 1999; Hari *et al*., 2021). Other than knowing the approximate location of the first order neurons involved in PAD evoked in Ia afferents (Jankowska *et al*., 1981), we know little about them, unlike the first order neurons involved in PAD evoked in LTMRs (CCK+ neurons, mentioned above). We can surmise that they may be propriospinal neurons to account for the widespread radiating nature of this PAD, where a single nerve stimulation can evoke PAD many segments away and across the midline (Barron & Matthews, 1938; Lucas-Osma *et al*., 2018). Further, these first order neurons likely express specialized persistent intrinsic currents that allows them to produce very long responses, since PAD far outlasts the brief sensory activation needed to trigger it (Barron & Matthews, 1938; Lucas-Osma *et al*., 2018).

The local spinal central pattern generator (CPG) circuits that produce walking or other complex movements like flexion withdrawal reflexes also strongly modulate PAD (Jankowska *et al*., 1965; Anden *et al*., 1966; Gossard, 1996; Rossignol *et al*., 2006), but again the details of the circuits involved remain uncertain, other than knowing that GABA_axo_ neurons also produce this PAD, and likely some propriospinal neuron population allows this PAD to also be widespread across many segment levels and associated muscles. We do however know a lot about the neurons in the CPG itself (Leblond *et al*., 2003; Kiehn, 2016; Gosgnach *et al*., 2017; Ziskind-Conhaim & Hochman, 2017). Of these CPG-related neurons, the V3 neuron is a good candidate to be involved in generating PAD, as it has extensive propriospinal and commissural axons (Zhang *et al*., 2008). Indeed, in the course of our recent studies of locomotion and spasticity in mice (Lin *et al*., 2019) we noticed that optogenetic activation V3 neurons not only activates motoneurons, but also produces a pronounced PAD (published in abstract form)(Li *et al*., 2015), and this finding is the subject of this paper. Previous unpublished work has suggested that V3 neurons receive sensory input (published in abstract form)(Deska-Gauthier *et al*., 2018), and thus we started here verifying this, which would allow these neurons to not only be involved in PAD during locomotion, but also more generally evoke PAD in response to sensory stimulation. We report here that V3 neurons are not only involved in PAD, but are essential for a large portion of sensory-evoked PAD in proprioceptive afferents, even in the absence of locomotion, suggesting that V3 neurons may be the illusive first order neuron on the proprioceptive trisynaptic PAD circuit. We also unexpectedly found that V3 neurons produce a strong PAD that does not depend on GABA, but instead is in part mediated by NMDA receptors. While such NMDA-dependent PAD has been observed before during stimulation of high threshold C or Aδ afferents (Russo *et al*., 2000; Zimmerman *et al*., 2019), its origin is unclear, and thus we also examined how V3 neurons contributed NMDA-related PAD. In summary, our find that V3 neurons are essential to PAD generation suggests that PAD circuits are not an isolated sensory control system, but a part of the overall sensorimotor system that controls movement, since V3 neurons have many motor functions, including directly driving motoneurons, switching on locomotor activity and coordinated interlimb activity (Zhang *et al*., 2008; Chopek *et al*., 2018; Lin *et al*., 2019; Bohm *et al*., 2022).

## METHODS

### Adult mice strains used

Recordings were made from V3 neurons, group Ia afferents, dorsal roots (DRs) and ventral roots in the sacrocaudal spinal cord of adult mice (4 – 6 months old, both female and male equally; strains detailed below). All experimental procedures were approved by the University of Alberta Animal Care and Use Committee, Health Sciences division. We evaluated V3 neurons in mice with Cre recombinase expressed under the Sim1 promotor region, as detailed previously (Zhang *et al*., 2008; Lin *et al*., 2019). These V3 neurons were visualized with Cre driven fluorophores (tdTom or EYFP), silenced with Cre driven VGLUT2 knockout (V3 neurons use VGLUT2 for vesicular glutamate transport), activated optogenetically using Cre driven channelrhodopsin-2 (ChR2) or inhibited optogenetically with Cre driven archaerhodopsin-3 (AchT).

The Sim1-Cre mice were obtained from two sources: 1) Sim1-Cre-ki mice, where Cre-recombinase is knocked in (ki) under the endogenous Sim1 promoter of the host genome (Zhang *et al*., 2008), and so expresses Cre under the control of Sim1 expression (obtained courtesy of Dr. M. Goulding, Salk Inst., USA), and 2) Sim1-Cre-tg mice, where an artificially generated Sim1-Cre transgene (tg) is inserted into the mouse genome (The Jackson Laboratory, Strain #:006395) and transgene expression is observed in most areas that endogenously express Sim1, including the sacral spinal cord that we study here. Results obtained from the sacral spinal cord with Sim1-Cre-ki and Sim1-Cre-tg were similar and combined, and hereafter both these mice were abbreviated Sim1-Cre mice.

The following floxed reporter strains were employed (Madisen *et al*., 2010; Madisen *et al*., 2012): 1) B6.Cg-*Gt(ROSA)26Sor_tm14(CAG-tdTomato)Hze_* and B6.Cg-*Gt(ROSA)26Sor_tm9(CAG-tdTomato)Hze_* mice (abbreviated R26_LSL-tdTom_ mice; The Jackson Laboratory, Stock # 007914 and #007909; tdTomato fluorescent protein expressed under the R26::CAG promotor in cells that co-express Cre), 2) B6J.129S6(129S4)-Slc17a6_tm1Lowl/RujfJ_ mice where the VGLUT2 gene (Exon 2) is flanked by loxP sites and Cre recombinase excises the VGLUT2 exon to generated a knockout (abbreviated VGLUT2_flox_ mice), 3) B6;129S-*Gt(ROSA)26Sor_tm32(CAG-COP4*H134R/EYFP)Hze_* mice (abbreviated R26_LSL-ChR2-EYFP_ mice; The Jackson Laboratory, Stock # 012569; ChR2-EYFP fusion protein expressed under the R26::CAG promotor in cells that co-express Cre because a loxP-flanked STOP cassette, LSL, prevents transcription of the downstream ChR2-EYFP gene), and 4) B6;129S-*Gt(ROSA)26Sor_tm35.1(CAG-aop3/GFP)Hze_* mice (abbreviated R26_LSL-Arch3-GFP_ mice; The Jackson Laboratory Stock # 012735; Arch3-GFP fusion protein expressed under the R26::CAG promotor in cells that co-express Cre). Offspring without the Sim1-cre or mutation, but with the effectors tdTom, VGLUT2_flox_, ChR2, or Arch3 were used as controls.

Sim1-cre mice were crossed with homozygous reporter strains to generate Sim1-cre_+_;R26_LSL-tdTom_, Sim1-cre;VGLUT2_flox_, Sim1-cre;R26_LSL-ChR2-EYFP_, and Sim1-cre; R26_LSL-Arch3-GFP_ mice that we abbreviate: Sim1//tdTom, Sim1//VGLUT2_KO_, Sim1//ChR2, and Sim1//Arch3 mice, respectively.

We also studied GABAergic neurons in mice with GFP expressed under GAD1 (GAD1-GFP, obtained courtesy of Dr. Peter Smith), and crossed these with Sim1//tdTom mice. Additionally we used mice with Cre inserted after GAD2, the latter as previously detailed (Hari *et al*., 2021), using *Gad2_tm1(cre/ERT2)Zjh_* mice (abbreviated Gad2_CreER_ mice; The Jackson Laboratory, Stock # 010702; CreER_T2_ fusion protein expressed under control of the endogenous *Gad2* promotor). From these we generated GAD2//ChR2 mice to optogenetically activate them as for Sim1//ChR2 mice detailed above.

### Chronic spinal cord injury (SCI)

Some mice were studied after a chronic S2 spinal transection, described previously (Lin *et al*., 2019). Briefly, adult mice were transected at the S2 sacral spinal level at ∼2 mo of age (adult mice), and recordings were made 1.5–3 mo after injury when their affected muscles became spastic as detailed previously. Under general anesthesia (100 mg/kg ip ketamine hydrochloride and 10 mg/kg ip xylazine) and using aseptic technique, the L2 lumbar vertebra was exposed and a laminectomy performed to expose the S2 sacral spinal cord. The dura was cut transversely, and ∼0.1 ml of Xylocaine (1%) was applied to the exposed spinal cord. With the use of a surgical microscope, the exposed pia was held with fine forceps and the spinal cord was transected by aspirating a short section of the spinal cord using a fine suction tip. Caution was needed to avoid damaging the anterior artery and dorsal vein because the sacrocaudal spinal cord dies without these vessels. The dura was closed with 8-0 silk sutures, and the muscle and skin were sutured in layers using 5-0 silk. Immediately following surgery, the mouse was placed in a recovery cage located on a heating blanket and allowed to recover.

### In vitro recording in whole adult spinal cords

Mice were anaesthetized with urethane (for mice 0.11 g/100 g, with a maximum dose of 0.065 g), a laminectomy was performed, and then the entire sacrocaudal spinal cord was rapidly removed and immersed in oxygenated modified artificial cerebrospinal fluid (mACSF), as detailed previously (Lin *et al*., 2019). Spinal roots were removed, except the sacral S2, S3, S4 and caudal Ca1 ventral and dorsal roots on both sides of the cord. After 1.5 hours in the dissection chamber (at 20° C), the cord was transferred to a recording chamber containing normal ACSF (nACSF) maintained at 23°C, with a flow rate > 3 ml/min. A one-hour period in nACSF was given to wash out the residual anaesthetic prior to recording, at which time the nACSF was recycled in a closed system. The cord was secured onto tissue paper at the bottom of a rubber (Silguard) chamber by insect pins in connective tissue and cut root fragments. The left side of the cord was usually oriented upwards when making intracellular recording from Ia afferents in the dorsal horn, whereas the cord was oriented with its left side upwards when making recordings from motoneurons or V3 neurons. The laser beam used for optogenetics was focused vertically downward on the V3 neurons or GAD2 neurons, as detailed below.

### Optogenetic regulation of V3 neurons

The Sim1//ChR2 or Sim1//Arch3 mice were used to optogenetically excite or inhibit V3 neurons (with 447 nm D442001FX and 532 nM LRS-0532-GFM-00200-01 lasers from Laserglow Technologies, Toronto), respectively, using methods we previously described (Lin *et al*., 2019; Hari *et al*., 2021). GAD2//ChR2 mice were likewise used to excite GABAergic neurons. Light was derived from the laser passed through a fibre optic cable and then a half cylindrical prism the length of about two spinal segments (8 mm; 3.9 mm focal length, Thor Labs, Newton, USA,), which collimated the light into a narrow long beam (200 µm wide and 8 mm long). This narrow beam was usually focused longitudinally on the left side of the spinal cord to target many V3 neurons at once. ChR2 rapidly depolarizes neurons (Zhang *et al*., 2011), and thus we used 5 – 10 ms light pulses to activate V3 neurons. Light was kept at a minimal intensity, 2 - 3x T, where T is the threshold to evoke a light response, which made local heating from light unlikely. Arch3 is a proton pump that is activated by green light, leading to a hyperpolarization and slowly increased pH (over seconds), both of which inhibit the neurons (Zhang *et al*., 2011; El-Gaby *et al*., 2016). Thus, we used longer light pulses to inhibit V3 neurons.

To directly confirm the presence of functional ChR2 expression in V3 neurons of Sim1//ChR2 mice we recorded from them with similar methods and intracellular electrodes that we used to record from afferents (see below). Electrodes were advanced into these cells through the lateral edge of the cord, and their identity established by a direct response to light activation of the ChR2 construct (5 – 10 ms light pulse, 447 nm), without a synaptic delay (<1 ms) and continued light response after blocking synaptic transmission.

### Dorsal stimulation

During intracellular recordings all dorsal roots were mounted on silver-silver chloride wires above the nASCF of the in vitro chamber and covered with grease (a 3:1 mixture of petroleum jelly and mineral oil) for monopolar stimulation (Lucas-Osma *et al*., 2018; Lin *et al*., 2019). This grease was surrounded by a more viscous synthetic high vacuum grease to prevent oil leaking into the bath flow. Bipolar stimulation was also used at times to reduce the stimulus artifact. DRs were stimulated with a constant current stimulator (Isoflex, Israel) with short pulses (0.1 ms). During grease-gap recording (detailed below) only the Ca1 DR was mounted for stimulation when we record PAD from other DRs (S2-S4 DRs).

### Intracellular recording and labelling sensory axons in the dorsal horn

Intracellular recordings were made from group Ia afferents with electrodes made from glass capillary tubes (1.5 mm and 0.86 mm outer and inner diameters, respectively; with filament; 603000 A-M Systems; Sequim, USA) pulled with a Sutter P-87 puller (Flaming-Brown; Sutter Instrument, Novato, USA), filled with either 1 M K-acetate and 1 M KCl or 500 mM KCl in 0.1 Trizma buffer with 5 - 10% neurobiotin; Vector Labs, Birmingame, USA), and beveled to 30 - 40 MΩ using a rotary beveller (Sutter BV-10)(Hari *et al*., 2021). Intracellular recording was performed with an Axoclamp2B amplifier (Axon Inst. and Molecular Devices, San Jose, USA). Recordings were low pass filtered at 10 kHz and sampled at 30 kHz (Clampex and Clampfit; Molecular Devices, San Jose, USA. Electrodes were advanced into afferents of the sacrocaudal spinal cord with a stepper motor (Model 2662, Kopf, USA, 10 µm steps at maximal speed, 4 mm/s), usually at the boundary between the dorsal columns and dorsal horn gray matter, where axons bundle together densely, as they branch and descend to the ventral horn. Prior to penetrating afferents, we recorded the extracellular (EC) afferent volley following dorsal root (DR) stimulation (0.1 ms pulses, 3xT, T: afferent volley threshold, where T = ∼3 uA, repeated at 1 Hz), to determine the minimum latency and threshold of afferents entering the spinal cord. The group Ia afferent volley occurs first with a latency of 0.5 - 1.0 ms, depending on the root length (which were kept as long as possible, 10 - 20 mm). Upon penetration, afferents were identified with direct orthodromic spikes evoked from DR stimulation. We focused on the lowest threshold proprioceptive group Ia afferents, identified by their direct response to DR stimulation, very low threshold (< 1.5 x T, T: afferent volley threshold), and short latency (group Ia latency, coincident with onset of afferent volley). Clean axon penetrations without injury occurred abruptly with the membrane potential settling rapidly to near – 70 mV, and > 70 mV spikes usually readily evoked by DR stimulation or brief current injection pulses (1 – 3 nA, 20 ms, 1 Hz). Sensory axons also had a characteristic >100 ms long depolarization following stimulation of a dorsal root (PAD) and short spike afterhyperpolarization (AHP ∼ 10 ms), which further distinguished them from other axons or neurons. Injured axons had higher resting potentials (> - 60 mV), poor spikes (< 60 mV) and low resistance (to current pulse; R_m_ < 10 MΩ), and were discarded.

Some of the proprioceptive afferents that we recorded intracellularly were subsequently filled with neurobiotin by passing a very large positive 2 - 4 nA current with 90% duty cycle (900 ms on, 100 ms off) for 10 - 20 min. The identity of group Ia proprioceptive afferents were then confirmed anatomically by their unique extensive innervation of motoneurons (Lucas-Osma *et al*., 2018). Prior to penetrating and filling axons with neurobiotin filled electrodes, a small negative holding current was maintained on the electrodes to avoid spilling neurobiotin outside axons.

### Dorsal and ventral root grease gap recording

In addition to recording directly from single proprioceptive axons, we employed a grease gap method to record the composite intracellular response of many sensory axons or motoneurons by recording from dorsal and ventral roots, respectively, as previously detailed (Hari *et al*., 2021). We mounted the freshly cut roots onto silver-silver chloride wires just above the bath, and covered them in grease over about a 2 mm length, as detailed above for monopolar stimulation. Return and ground wires were in the bath and likewise made of silver-silver chloride. Specifically for sensory axons, we recorded from the central ends of dorsal roots (S2-S4) cut within about 2 - 4 mm of their entry into the spinal cord, to give the compound potential from all afferents in the root (dorsal roots potential, DRP), which has previously been shown to correspond to PAD, though it is attenuated compared to the intracellular recordings of PAD (Lucas-Osma *et al*., 2018). For optogenetic experiments we additionally added silicon carbide powder (9 % wt, Tech-Met, Markham) to the grease to make it opaque to light and minimize light induced artifactual current in the silver-silver chloride recording wire during optogenetic activation of ChR2 (detailed below). Likewise, we covered our bath ground and recording return wires with a plastic shield to prevent stray light artifacts. The dorsal root recordings were amplified (2,000 times), high-pass filtered at 0.1 Hz to remove drift, low-pass filtered at 10 kHz, and sampled at 30 kHz (Axoscope 8; Axon Instruments/Molecular Devices, Burlingame, CA).

The composite EPSPs in many motoneurons were likewise recorded from the central cut end of ventral roots (S3-S4) mounted in grease (grease gap), which has also previously been shown to yield reliable estimates of the EPSPs, though again attenuated by the distance from the motoneurons (Hari *et al*., 2021). The monosynaptic EPSPs were again identified as monosynaptic by their rapid onset (first component, ∼1 ms after afferent volley arrives in the ventral horn; see below), lack of variability in latency (< 1 ms jitter), persistence at high rates (10 Hz) and appearance in isolation at the threshold (T) for evoking EPSPs with DR stimulation (< 1.1xT, T ∼ afferent volley threshold), unlike polysynaptic reflexes which varying in latency, disappear at high rates, and mostly need stronger DR stimulation to activate. Usually the Ca1 or S4 roots were stimulated to evoke the EPSPs.

### Drugs and solutions

Two kinds of artificial cerebrospinal fluid (ACSF) were used in these experiments: a modified ACSF (mACSF) in the dissection chamber prior to recording and a normal ACSF (nACSF) in the recording chamber. The mACSF was composed of (in mM) 118 NaCl, 24 NaHCO3, 1.5 CaCl2, 3 KCl, 5 MgCl2, 1.4 NaH2PO4, 1.3 MgSO4, 25 D-glucose, and 1 kynurenic acid. Normal ACSF was composed of (in mM) 122 NaCl, 24 NaHCO3, 2.5 CaCl2, 3 KCl, 1 MgCl2, and 12 D-glucose. Both types of ACSF were saturated with 95% O2-5% CO2 and maintained at pH 7.4. The drugs sometimes added to the ACSF were APV (NMDA receptor antagonist), CNQX (AMPA antagonist), gabazine (GABA_A_ antagonist), and TTX (TTX-citrate; Toronto Research Chemicals, Toronto). Drugs were first dissolved as a 10 - 50 mM stock in water or DMSO before final dilution in ACSF.

### Immunohistochemisty

Sim1//tdTom or Sim1//ChR2-EYFP mice were euthanized with Euthanyl (BimedaMTC; 700 mg/kg) and perfused intracardially with 10 ml of saline for 3 – 4 min, followed by 40 ml of 4% paraformaldehyde (PFA; in 0.1 M phosphate buffer at room temperature), over 15 min (Gabra5-KO mice also fixed similarly). Then spinal cords of these mice were post-fixed in PFA for 1 hr at 4°C, and then cryoprotected in 30% sucrose in phosphate buffer (∼48 hrs). Alternatively, spinal cords where sensory axons were injected with neurobiotin in vitro were left in the recording chamber in oxygenated nACSF for an additional 4 – 6 hr to allow time for diffusion of the neurobiotin throughout the axon and then the spinal cord was immersed in 4% paraformaldehyde (PFA; in phosphate buffer) for 20-22 hours at 4°C, cryoprotected in 30% sucrose in phosphate buffer for 24-48 hours. Following cryoprotection all cords were embedded in OCT (Sakura Finetek, Torrance, CA, USA), frozen at −60C with 2-methylbutane, cut on a cryostat NX70 (Fisher Scientific) in sagittal or transverse 25 µm sections, and mounted on slides. Slides were frozen until further use.

The tissue sections on slides were first rinsed with phosphate buffered saline (PBS, 100 mM, 10 min) and then again with PBS containing 0.3% Triton X-100 (PBS-TX, 10 min rinses used for all PBS-TX rinses). Next, for all tissue, nonspecific binding was blocked with a 1 h incubation in PBS-TX with 10% normal goat serum (NGS; S-1000, Vector Laboratories, Burlingame, USA) or normal donkey serum (NDS; ab7475, Abcam, Cambridge, UK). Sections were then incubated for at least 20 hours at room temperature with a combination of the following primary antibodies in PBS-TX with 2% NGS or NDS: guinea pig anti-VGLUT1 (1:1000; AB5905, Sigma-Aldrich, St. Louis, USA), guinea pig anti-VGLUT2 (1:20,000, AB225-I, Sigma-Aldrich, St. Louis, USA), guinea pig anti-GAD2/GAD65 (1:500; 198 104; Synaptic Systems); chicken anti-VGAT (1:500; 131 006, Synaptic Systems, Goettingen, Germany), rabbit anti-VGAT (1:500; AB5062P, Sigma-Aldrich, St. Louis, USA), rabbit anti-EYFP (1:500; orb256069, Biorbyt, Riverside, UK), mouse anti-bassoon (1:400, ENZO SAP7F407, MJS Biolynx Inc), goat anti-RFP (1:500; orb334992, Biorbyt, Riverside, UK), rabbit anti-RFP (1:500; PM005, MBL International, Woburn, USA), and rabbit anti-GFP (1:500, A11122, ThermoFisher Scientific, Waltham, USA). Genetically expressed EYFP (labelled by GFP-ab), tdTom (labelled by RFP-ab) and GFP were amplified with the above antibodies, rather than rely on the endogenous fluorescence. When anti-mouse antibodies were applied in mice tissue, the M.O.M (Mouse on Mouse) immunodetection kit was used (M.O.M; BMK-2201, Vector Laboratories, Burlingame, USA) prior to applying antibodies. This process included 1h incubation with a mouse Ig blocking reagent. Primary and secondary antibody solutions were diluted in a specific M.O.M diluent.

The following day, tissue was rinsed with PBS-TX (3x 10 min) and incubated with fluorescent secondary antibodies. The secondary antibodies used included: goat anti-rabbit Alexa Fluor 555 (1:200; A32732, ThermoFisher Scientific, Waltham, USA), goat anti-rabbit Alexa Fluor 647 (1:500, ab150079, Abcam, Cambridge, UK), goat ant-rabbit Pacific orange (1:500; P31584, ThermoFisher Scientific, Waltham, USA), goat anti-mouse Alexa Fluor 647 (1:500; A21235, ThermoFisher Scientific, Waltham, USA), goat anti-mouse Alexa Fluor 488 (1:500; A11001, ThermoFisher Scientific, Waltham, USA), goat anti-mouse Alexa Fluor 555 (1:500; A28180, ThermoFisher Scientific, Waltham, USA), goat anti-guinea pig Alexa Fluor 647 (1:500; A21450, ThermoFisher Scientific, Waltham, USA), goat anti-chicken Alexa Fluor 405 (1:200; ab175674, Abcam, Cambridge, UK), goat anti-chicken Alexa Fluor 647 (1:500; A21449, ThermoFisher Scientific, Waltham, USA), donkey anti-goat Alexa Fluor 555 (1:500; ab150130, Abcam, Cambridge, UK), donkey anti-rabbit Alexa Fluor 488 (1:500; A21206, ThermoFisher Scientific, Waltham, USA), Streptavidin-conjugated Alexa Fluor 488 (1:200; 016-540-084, Jackson immunoResearch, West Grove, USA) or Streptavidin-conjugated Cyanine Cy5 (1:200; 016-170-084, Jackson immunoResearch, West Grove, USA) in PBS-TX with 2% NGS or NDS, applied on slides for 2 h at room temperature. The latter streptavidin antibodies were used to label neurobiotin filled afferents. After rinsing with PBS-TX (2 times x 10 min/each) and PBS (2 times x 10 min/each), the slides were covered with Fluoromount-G (00-4958-02, ThermoFisher Scientific, Waltham, USA) and coverslips (#1.5, 0.175 mm, 12-544-E; Fisher Scientific, Pittsburg, USA).

Standard negative controls in which the primary antibody was either 1) omitted or 2) blocked with its antigen (quenching) were used to confirm the selectivity of the antibody staining, and no specific staining was observed in these controls. Previous tests detailed by the manufactures further demonstrate the antibody specificity, including quenching, immunoblots (Western blots), co-immunoprecipitation, and/or receptor knockout.

Image acquisition was performed by confocal microscopy (Leica TCS SP8 Confocal System). All the confocal images were taken with a 63x (1.4 NA) oil immersion objective lens or a 20x water emersion lens and 0.1 µm optical sections that were collected into a z-stack over 10–20 µm. Excitation and recording wavelengths were set to optimize the selectivity of imaging the fluorescent secondary antibodies. Large areas were imaged with the Tilescan option in Leica Application Suite X software (Leica Microsystems CMS GmbH, Germany).

### Statistical analysis

Data were analyzed in Clampfit 8.0 (Axon Instruments, USA) and Sigmaplot (Systat Software, USA). A Student’s *t*-test or ANOVA (as appropriate) was used to test for statistical differences between variables, with a significance level of *P <* 0.05 (two tailed). Power of tests was computed with α = 0.05 to design experiments. A Kolmogorov-Smirnov test for normality was applied to the data set, with a *P <* 0.05 level set for significance. Most data sets were found to be normally distributed, as is required for a *t*-test. For those that were not normal a Wilcoxon Signed Rank Test was instead used with *P <* 0.05. Effects in male and female animals were similar and grouped together in analysis. Axons and motoneurons were recorded in vitro from widely separated locations (one segment apart or contralateral) within the whole spinal cord, and are considered independent; so statistics were performed across all neurons (*n*) from all animals, though comparing across animal averages also showed significant changes, with 3 or more animals used per condition. Data are indicated by mean ± standard deviation.

## RESULTS

We began by genetically labelling V3 neurons in Sim1-cre//tdTom mice (Fig 1A) or Sim1-cre//ChR2-EYFP mice (not shown) to examine the role of V3 neurons in the putative trisynaptic circuit underlying sensory-evoked PAD (Fig 1). These neurons were extensively innervated with sensory afferent terminals labelled with VGLUT1 (Fig 1A-B)(Todd *et al*., 2003), including terminals from group Ia afferents filled with neurobiotin (Fig 1B). Overall, 91% of V3 neuron somas we examined were innervated by afferents (VGLUT1_+_; n = 21/23 neurons from intermediate zone). Furthermore, in the sacrocaudal spinal cord that we studied the V3 neurons and their extensive processes were predominantly located at intermediate spinal levels near the central canal (∼laminae 6, 7 and 8, as well as the sacral dorsal commissure), as previously detailed for the sacral cord (Borowska *et al*., 2013), well positioned to interact with nearby GABAergic neurons that mediate PAD (detailed later)(Hari *et al*., 2021).

**Fig. 1.**
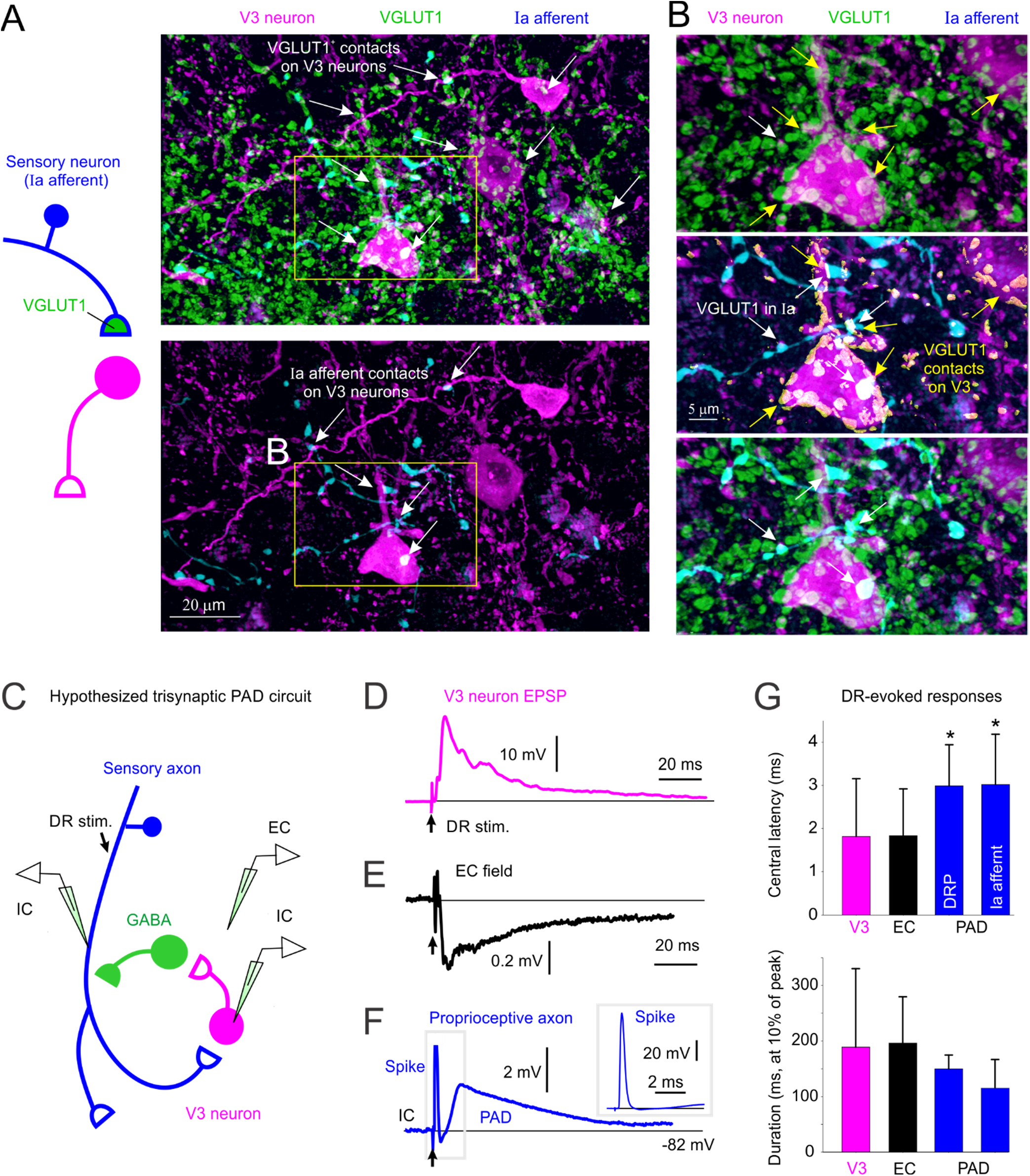
Monosynaptic connections from sensory afferents to V3 neurons. *A*: V3 neurons in the spinal cord intermediate laminae decorated with VGLUT1_+_ contacts from afferents, some of which are Ia afferent contacts labelled by an intracellular injection of neurobiotin (blue). Spinal cord from Sim1//tdTom mouse. *B*: Expanded image from *A* with 3D reconstructed VGLUT1_+_ afferent contacts on V3 neurons labelled in yellow. *C*: Schematic of putative trisynaptic circuit where Ia afferents (blue) innervate V3 neurons, which in turn innervate GABAergic neurons. which return to innervate Ia afferents, ultimately producing PAD. Experimental setup indicated where Ia afferents are activated by dorsal root (DR) stimulation (0.1 ms pulse, 2xT, sensory threshold) and response recorded with a sharp intracellular electrode in or near V3 neurons or afferents. *D*: monosynaptic EPSP recorded intracellularly in V3 neuron in response to dorsal roots stimulation (average of 10 trials at 0.3 Hz). *E*: Same V3 EPSP as in *D*, but population response recorded nearby, extracellulary to V3 neurons (EC field). *F*: Intra-axonal recording from Ia afferent during same DR stimulation, where the afferent is directly activated, as evident by an orthodromic spike, and following this a primary afferent depolarization (PAD) arises at a polysynaptic latency consistent with a circuit like in *C*. *G*: Central latencies and durations of V3 EPSPs, EC fields and PAD, where that latter was recorded either intracellularly in Ia afferents as in *F* (though in afferents of not directly activated by the DR stimulation, so the PAD onset can be judged) or by grease gap recordings from DR (dorsal root potential, DRP). Central latency measured relative to arrival time of the orthodromic spike at the spinal cord, as in F, though measured from the extracellular afferent volley (not shown, though see Lucas-Osma et al. 2018). V3 EPSP latencies were minimally near 1 ms (0.8 – 3 ms), which is monosynaptic since the in vitro adult spinal cord has a synaptic delay of ∼1 ms at 23_o_C. PAD latencies were 2 – 4 ms, with an average of 3 ms, consistent with the trisynaptic circuit of C, though possibly with also some disynaptic innervation. *, significantly longer latency of PAD (Ia PAD or DRP) compared to V3 EPSP or its EC field latency, *P* < 0.05, *n* = 6 V3 neurons, 6 EC fields, 44 DRPs and 5 Ia afferents.

To examine the direct action of this sensory innervation we made intracellular recordings from V3 neurons (Fig 1C-G). This revealed large, long-lasting EPSPs (V3 EPSPs) in response to low threshold sensory stimulation of the dorsal roots (DR, 1.5xT), with a rapid onset latency (Fig 1G), consistent with a minimally monosynaptic innervation from large proprioceptive or cutaneous afferents, as suggested by our confocal imaging (Fig 1A; the in vitro adult spinal cord has a synaptic latency of about 0.8 – 1 ms near room temperature)(Hari *et al*., 2021). Just outside V3 neurons a pronounced extracellular field (EC V3 field, negative, Fig 1E) was observed with a similar duration and monosynaptic latency as the V3 EPSP (Fig 1G), suggesting large numbers of V3 neurons were activated coincidently by the sensory innervation.

These long lasting V3 EPSPs were similar in duration and shape to depolarization of primary sensory afferents evoked by the same DR stimulation (sensory-evoked PAD), either measured directly by intracellular recording from group Ia afferents or indirectly by grease gap recording from the DRs, the latter which gives the population response of many large afferents (DRP, Fig 1F-G). When the same DR was stimulated that contained the recorded Ia afferent (Fig 1C) there was a direct orthodromic spike evoked if the stimulus intensity was above the afferent threshold, and this was followed by PAD (homonymous PAD), as depicted in Fig 1C, E. Such direct spikes compose the afferent volley that drives the V3 neurons monosynaptically in Fig 1D. Alternatively, when we stimulated an adjacent DR so there was no orthodromic spike, a similar PAD arose (heteronymous PAD, not shown, see Lucas et al. 2018). Either way, the sensory-evoked PAD activated by stimulating any DR (abbreviated drPAD) had a similar duration to the V3 response to the same DR stimulation (V3 EPSP; Fig 1G). As previously reported, the central latency of drPAD was about 2 - 4 ms (Fig 1F)(Lucas-Osma *et al*., 2018), longer than the V3 EPSP latency, and consistent with the classical assumption that PAD arises from a trisynaptic loop, such as in Fig 1C. This provides sufficient time for the V3 neurons to contribute to PAD, since their V3 EPSP has a central latency of half the PAD latency, as mentioned above, even taking into account possible delays in spike initiation in V3 neurons (0.5 – 4 ms; not shown).

In order for V3 neurons to be part of the trisynaptic drPAD circuit they need to directly contact GABAergic neurons (Fig 1C). To establish this, we crossed Sim1//tdTom mice with GAD1-GFP mice to label both V3 and GABAergic neurons, that latter including GAD2_+_ neurons which are the subset of GAD1_+_ neurons that form axoaxonic connections on afferents (Fig 2)(Betley *et al*., 2009). As expected, we found that V3 neurons extensively innervated GABAergic neurons, with presynaptic VGLUT2_+_ and Bassoon_+_ in these glutamateric V3 neurons, and postsynaptic GAD2 expression in GABAergic neurons labelled in GAD-GFP mice (Fig 2A-C). The majority of these V3 contacts were on GABAergic neurons in the intermediated and dorsal laminae (Fig 2E). Presynaptic bassoon was also expressed in the GABAergic neuron boutons near GAD2 and the V3 contacts onto these neurons, showing that the GABAergic neurons release GABA very near to where they receive V3 neuron innervation (Fig 2C), likely on a dendrite, a microcircuit arrangement that is common in GABAergic neurons (Russo *et al*., 2000; Shreckengost *et al*., 2010; Lucas-Osma *et al*., 2018). This suggests that GABAergic neuron may also innervate V3 neurons, an opposite arrangement to what we expected. Indeed, we found many GABAergic (GAD_+_) contacts onto V3 neurons, though these were more dorsally located, than the V3 neuron contacts on GABAergic neurons (Fig 2D, E). Taken together, our anatomical results demonstrate that V3 neurons are part of the PAD circuit receiving sensory input and innervating GABAergic GAD2_+_ neurons that are known to in turn innervate sensory afferents (Fig 1C). However, this PAD circuit is in part inhibited by GABAergic input (Fig 2F), complicating the interpretation of the action of GABA receptor antagonists commonly used to study PAD, which would disinhibit the V3 neurons in the PAD circuit and spinal cord in general, as detailed below.

**Fig. 2.**
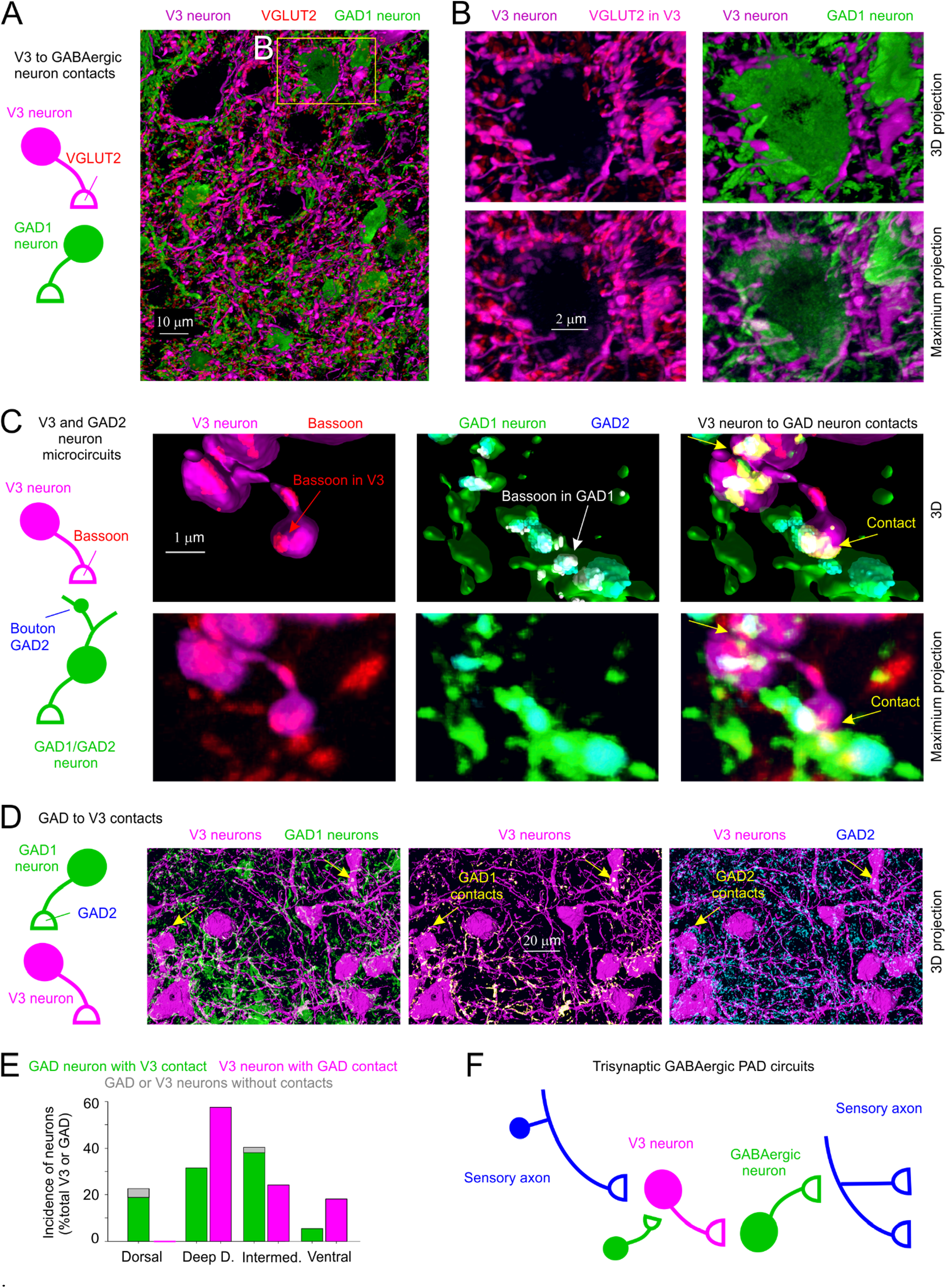
Connections between V3 neurons and GABAergic neurons that produce PAD. *A*-*B*: V3 neurons (tdTom) contacts onto GABAergic neurons, where presynaptic V3 neuron contacts are VGLUT2_+_(red), the vesicular transporter expressed in these glutamatergic V3 neurons. Spinal cord from mouse expressing Sim1//tdTom (magenta, V3) and GAD1-GFP (green, GABAergic neurons). *C*: Close up of V3 neuron contacts (Sim1//tdTom) onto GABAergic neuron (GAD1-GFP)), with V3 presynaptic terminal labelled with bassoon and GABAergic neuron labelled with GAD2, the latter to show that it is a GAD2_+_ neuron which uniquely innervates afferents (Todd refs). Bassoon is also expressed in the GABAergic neuron boutons near the GAD2 clusters and the V3 contacts. *D*: GABAergic (GAD1-GFP) neurons also innervate V3 neurons, with GAD2_+_ presynaptic contacts. *E*: Distribution and incidence of contacts between V3 neurons and GABAergic neurons in the dorsal, deep dorsal, intermediate and ventral laminae. *F*: Schematic summarizing trisynaptic circuit mediating PAD with the addition of a contact from GABAergic neurons onto V3 neurons that inhibits the circuit.

To quantify how important V3 neurons are drPAD evoked by proprioceptive sensory stimulation we used two approaches to block their function. First, we selectively knocked out (KO) glutamate function in V3 neurons in Sim1-cre// VGLUT2-flox mice (Sim1//VGLUT2_KO_), and second, we inhibited V3 neurons optogenetically by applying light to Sim1//ArchT mice, where the ArchT construct selectively expressed in V3 neurons hyperpolarizes these neurons and inhibits a proton pump (Fig 3). Both methods led to a substantial reduction drPAD evoked by stimulation of proprioceptors in the DR, which has previously shown to be largely mediated by GABAA receptors in this preparation (GABA PAD, detailed later)(Lucas-Osma *et al*., 2018). The ArchT silencing of V3 neurons was likely incomplete, but allowed within animal comparisons of changes in drPAD with light, yielding about a 50% reduction in drPAD (Fig 3A-C), suggesting that V3 neurons are responsible for at least half the PAD. The KO silencing of V3 neurons was likely complete (Fig 3D-E), but can only be compared across animals where drPAD variability was large, depending on the root size and quality of grease seal, but nevertheless on average led to an even larger reduction in drPAD than with ArchT. Importantly, with both these ArchT and KO experiments we evoked the PAD with very low threshold DR stimuli to ensure that only large proprioceptive Ia afferents were activated, which has previously been shown to mostly evoke PAD in proprioceptive afferents themselves, and not higher threshold cutaneous afferents (Jankowska *et al*., 1981; Lidierth & Wall, 1998; Rudomin & Schmidt, 1999; Zimmerman *et al*., 2019; Lalonde & Bui, 2020; Hari *et al*., 2021). Together these results indicate that the majority of sensory-evoked PAD in proprioceptive afferents is mediated through V3 neurons that act as the first order neurons (Fig 1C).

**Fig 3.**
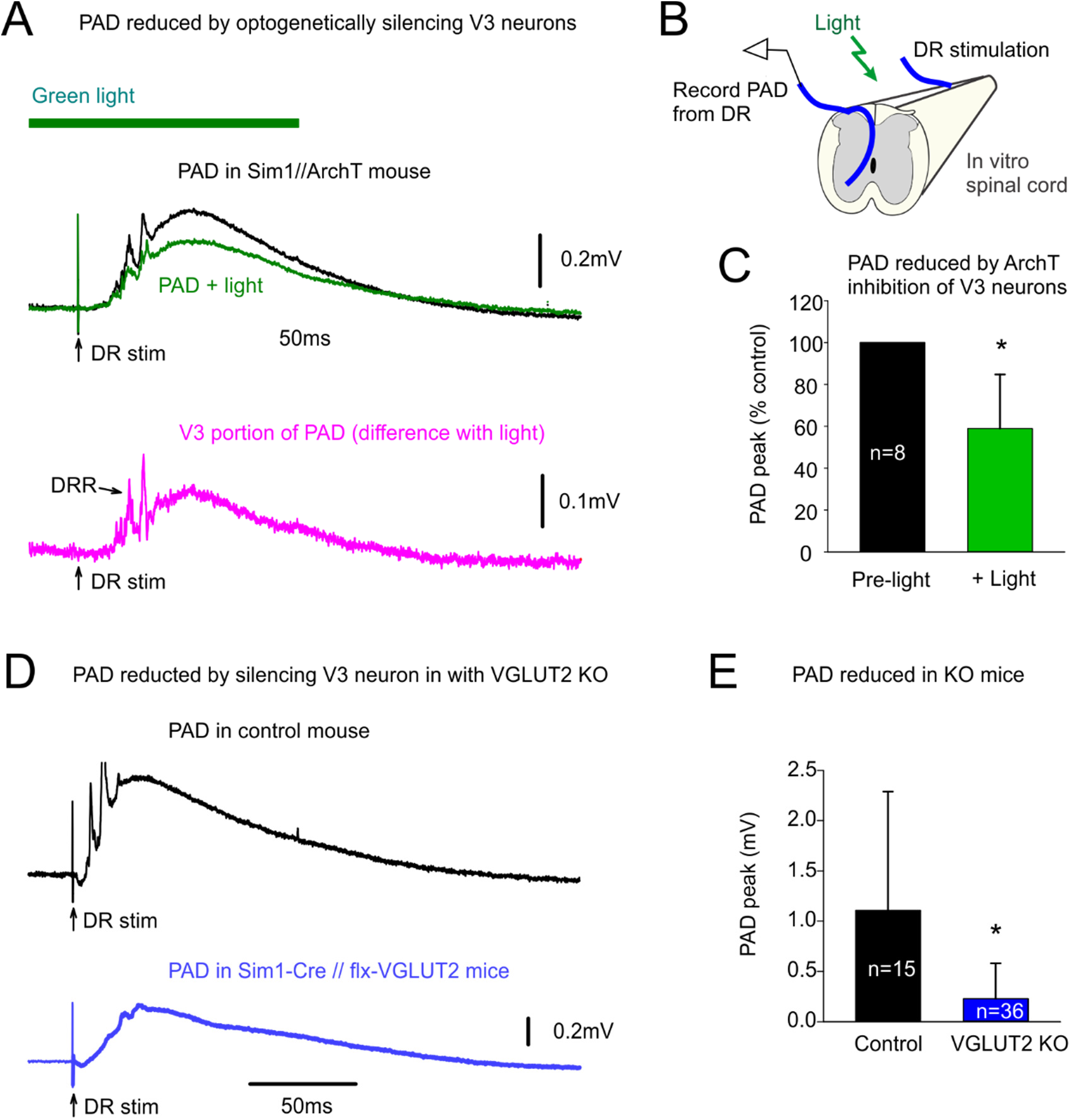
V3 neurons are essential for a large fraction of the PAD evoked by proprioceptive stimulation (drPAD). *A-B*: Typical PAD evoked by proprioceptive sensory stimulation of a DR (S4 DR, 0.1 ms, 1.1xT, T sensory threshold) in adjacent DR (DRP measured in S3 DR) in Sim1//ArchT mice, before and after silencing V3 neurons with light (532nm, 5 mW/mm_2_). Averages of 9 trials at 0.1 Hz. Light had no effect on drPAD in mice without ArchT; not shown, but as detailed previously (Hari, 2021). Lower trace: V3 dependent portion of drPAD including dorsal root reflexes (DRR), estimated from the change with light. *C*: Group averages of drPAD evoked by selective proprioceptive DR stimulation (1.1-1.3xT), * significant reduction with light, *P* < 0.05, *n* = 8. *D*: Typical drPAD evoked by proprioceptive stimulation (DR 2xT) as in *A*, but in mice without and with V3 neurons silenced by VGLUT2 KO; top: control mouse (Sim1-Cre lacking flx-VGLUT1); bottom: Sim1-Cre//flx-VGLUT2 mouse. *E*: Group averages of drPAD, * significantly smaller with KO, *P* < 0.05, *n* = 15 and 36 for control and KO mice respectively.

Considering that V3 neurons are part of the sensory PAD circuit, specifically innervating GABAergic neurons, we next examined Sim1//ChR2 mice where V3 neurons contained ChR2 so we could directly activate V3 neurons with a light pulse to examine their intrinsic properties and action on PAD. As mentioned above, V3 neurons in the sacral spinal cord were mostly located in the intermediate lamina with extensive arbors (Fig 4A). Thus, we recorded from neurons this region (as in Fig 4B), with V3 neurons penetrated through the lateral edge of the cord and identified by direct (non-delayed) responses to light activation of ChR2. Just like the long V3 EPSP observed with a brief DR stimulation (Figs 1D and 4C), a brief light pulse evoked a long-lasting response in V3 neurons (Fig 4D). These light responses were so large and characteristically long that they were also readily observed with extracellular recording (EC V3 field, seen as negative deflection). Individual trials in both the light and DR-evoked V3 responses showed considerable long-latency synaptic events (Fig 4C-D), and so we initially thought that perhaps the long-lasting nature of these V3 responses were due to reverberating synaptic circuits. While these synaptic events were eliminated by a complete block of all fast synaptic transmission (with CNQX, APV, gabazine and strychnine), this unexpectedly did not eliminate the long lasting V3 response to light, not decreasing the amplitude or duration (Fig 4D-E, G-I). However, subsequent application of TTX markedly reduced the light response amplitude and duration (Fig 4F-I), indicating that intrinsic sodium mediated persistent inward currents (Na PICs) contributed to the long lasting V3 responses. This Na PIC likely involves Nav1.8 and Nav 1.9 channels, since blockade required 2 – 4 times more TTX than needed to block axon conduction and these channels are known to be somewhat TTX-resistant (Russo *et al*., 2000; de Lera Ruiz & Kraus, 2015).

**Fig 4.**
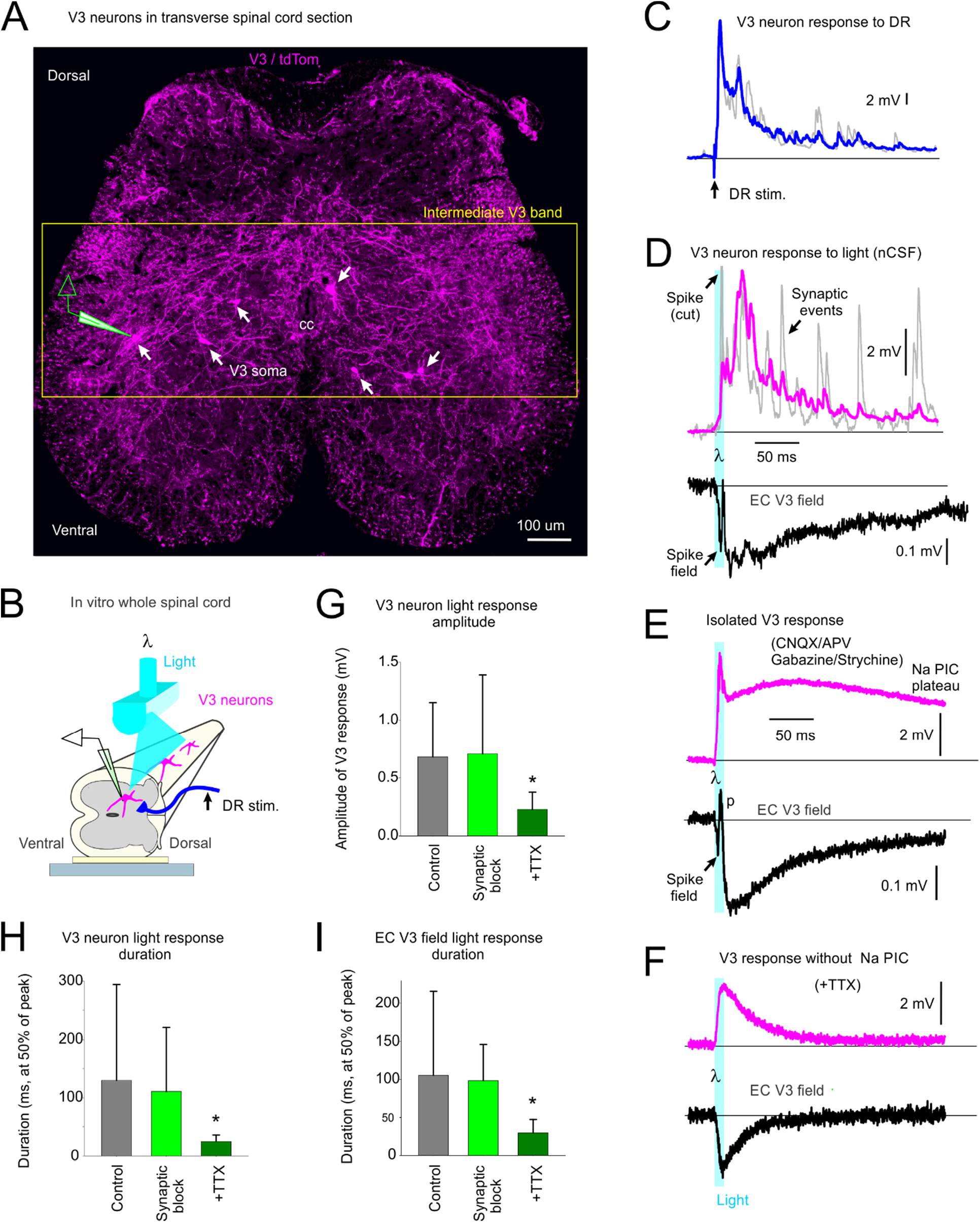
Long-lasting response evoked inV3 neurons by brief activation, mediated by persistent sodium currents. *A*: Transverse section of sacral S3 spinal cord of Sim1//tdTom mice showing V3 neurons with their extensive arborizations throughout much of the intermediate lamina, where we recorded from V3 neurons intracellularly. Repeated image from Fig 7, where details given. *B*: Arrangement for intracellular or extracellular recording from V3 neurons in Sim1//ChR2 mice, with neurons identified by a direct response to a light pulse (10 ms pulse, λ = 447nm laser, 0.7 mW/mm_2_, 3xT, T light threshold to evoke response, laser columnated and aligned axially along multiple segments), with no synaptic delay. *C*: Long-lasting V3 neuron EPSP in response to DR stimulation (0.1 ms, 2xT; as in Fig 1D), with blue line average of 10 trials (at 0.3 Hz), and grey line individual trial showing many synaptic events. *D*: Long-lasting V3 neuron response to the light pulse (10 ms, 1.5xT), with magenta line average of 10 trials, and grey line individual trial showing many fast synaptic events superimposed, as well as a fast spike at the onset of the response. Lower black line is the extracellular recording (EC field, averages of 10 trials) of the population response of V3 neurons to the same light pulse, which also showed a fast spike. *E*: Same as *D*, but after blocking synaptic transmission (with 50 µM CNQX, 50 µM APV, 50 µM gabazine and 5 µM strychnine), showing a long-lasting V3 neuron depolarization (plateau potential) from voltage-gated currents intrinsic to the V3 neurons. *F*: Same as *E*, but after also blocking sodium channels with TTX (4 µM) which eliminated the long-lasting response (and fast spike), showing that the plateau potential in *E* was due to persistent sodium currents (Na PICs). *G-I*: Group averages of V3 neuron responses to the brief light pulse, with amplitudes (quantified at 60 ms latency) and durations shown in control conditions, after blocking synaptic transmission, and then after also applying TTX. Duration measured at 50% peak amplitude. * significant reduction with TTX but not synaptic blockade, P < 0.05, n = 11 for intracellular V3 recordings, and n = 16 for EC V3 fields.

The intracellular and extracellular V3 responses to light consistently exhibited fast synchronous spikes that arose at about a 2 – 5 ms latency, peaked at about 5 – 6 ms, and were unaffected by blocking synaptic transmission (Fig 4G-H). This suggests that V3-evoked responses in postsynaptic neurons should have a minimally 2 ms latency, with on average a 5 – 6 ms latency, even with monosynaptic connections, which makes latencies of PAD-evoked by light slower than those evoked by DR stimulation, as detailed below. Likely, ChR2 is less effective at depolarizing V3 neurons than natural synaptic input, accounting for its slower action (as we have also seen in other neurons, Hari et al. 2021), and so caution must be used in interpreting latencies of ChR2 responses.

Interestingly, the light-evoked fast synchronous spikes recorded intracellularly in V3 neurons of Sim1//ChR2 mice were sometimes not full height (Fig 4E), suggesting that they arose somewhere in the long propriospinal axons of V3 neurons but did not always fully propagate back to the soma where we usually recorded, and so we observed only its passively attenuated spike (failure potential)(Hari *et al*., 2021). Indeed, the extracellular recording of this fast synchronous V3 evoked spike often had a large positive component, rather than the usual negative component expected for an extracellular spike arising near an extracellular electrode (Fig 4E, positive field p)(Hari *et al*., 2021). Again, this indicates that the spike arose far away in a portion of the V3 neuron’s axon illuminated by the light, but did not always actively propagate to the recording site, causing only a passive outward current near the electrode (a current sink rather than source of spike), previously detailed (Hari *et al*., 2021).

Considering that we typically applied the light mostly at and above (rostral) to the electrode during intracellular recording (for practical reasons with getting the electrode and laser in place), and that V3 neurons in the sacral cord tend to have a predominantly ascending propriospinal axons (as we detail below), we suggest that the light evoked spikes in many ascending propriospinal tracts and these spikes did not propagate antidromically to their parent cell bodies more caudally where the electrode was placed, which is not unusual because spike propagation failure is theoretically most likely when travelling from small distal branches to larger higher conductance branches (Goldstein & Rall, 1974).

To directly observe the action of V3 neurons on PAD we returned to recording from DR afferents in Sim1//ChR2 mice (Fig 5). As expected, a brief light activation of V3 neurons produced a long-lasting depolarization of afferents (termed V3 PAD; Fig 5A). This V3 PAD was similar to sensory-evoked PAD (drPAD) in that it had a long duration, large amplitude and short latency (Fig 5C-E; PAD estimated from the DRP with grease gap methods, which is about 10% of actual PAD size; see details in Lucas-Osma et al. 2018). Also, V3 PAD and drPAD were both sensitive to the GABAA receptor antagonist gabazine, consistent with both being in large part mediated by GABAA receptors (Fig 5A-B). V3 PAD had a slightly shorter latency (2.5 ms) compared to DR PAD (3 ms), consistent with its putative position two rather than three synapses away from the afferents in the trisynaptic circuit (Fig 5H). However, considering that spikes arise in V3 neurons slower with light compared to DR activation (right of Fig 5E and see above), these results suggest that the V3 neurons may also have an even more direct action on afferents (monosynaptic). Further, unlike with DR PAD, there were often two clearly distinguishable events in V3 PAD: one fast event with peak at about 10 ms (early peak) and a second with a much later peak at about 60 ms (late peak; Fig 5A, F, G). The sensory-evoked PAD for these same afferents had a single peak at about 30 ms, between the early and late V3 PAD peaks (Fig 5A, G).

**Fig 5.**
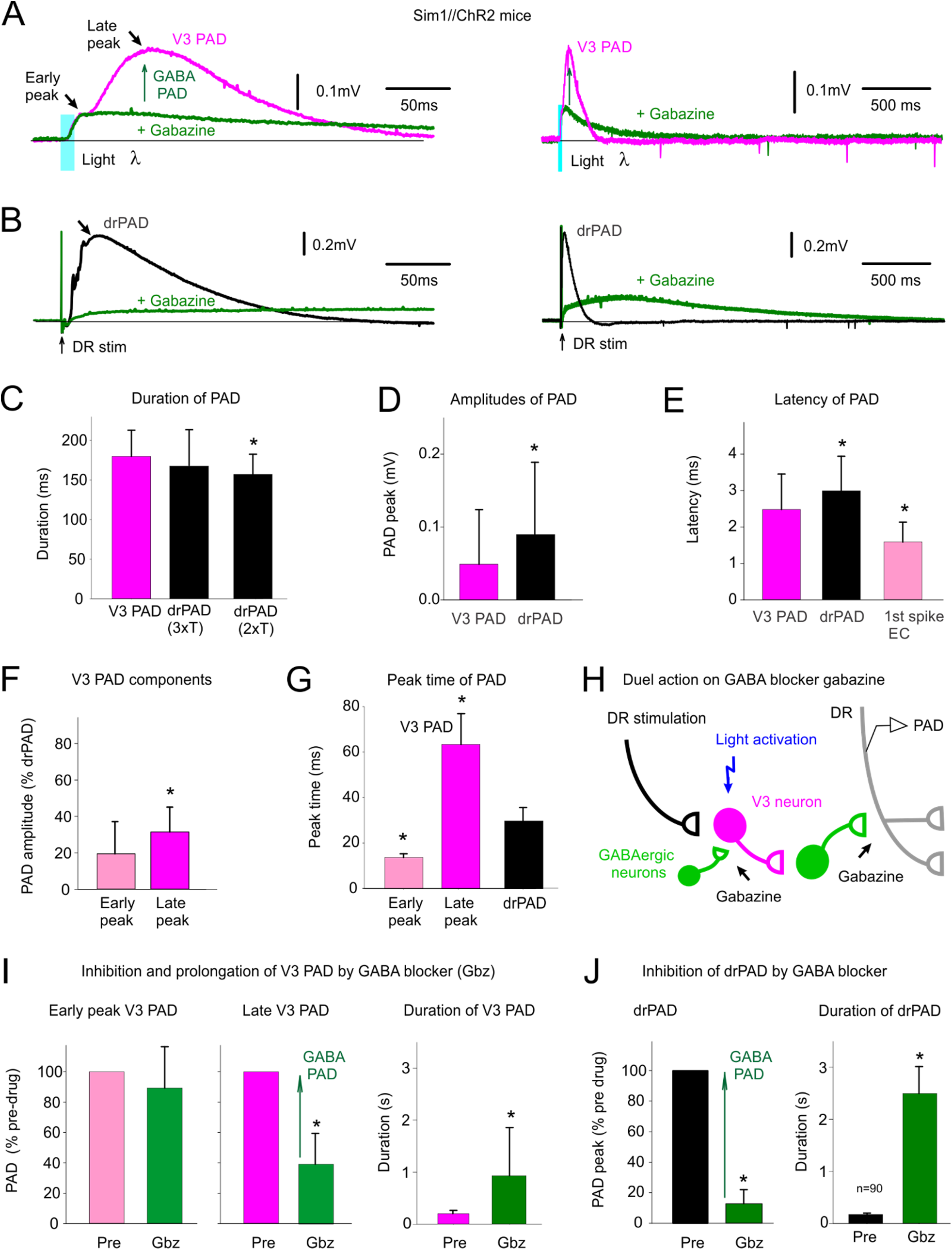
V3 neurons evoke a PAD (V3 PAD), partly mediated by GABA_A_ receptors. *A*: Brief optogenetic activation of V3 neurons in Sim1//ChR2 mice (10 ms pulse, λ = 447nm laser, 0.7 mW/mm_2_, 3xT, columnated laser light aligned axially to maximally activate V3 neurons over multiple segments, as in Fig 4B) evokes a PAD (termed V3 PAD) recorded in DR (S3 DR, DRP recorded with grease gap method), with an early peak and a late peak, as indicated. Blocking GABA_A_ receptors with gabazine (50 µM) blocks the late V3 PAD, but reveals a very long lasting PAD that starts at the early peak, as shown on the expanded time scale on the right. *B*: Similar to *A*, but brief DR stimulation evokes a drPAD that is mostly blocked by gabazine, but again reveals a very long lasting PAD. *C*: Group averages of durations of V3 PAD and drPAD evoked by 2xT and 3xT DR stimulation, prior to gabazine. * drPAD significantly less than V3 PAD, *n* = 28 V3 PADs and 50 drPADs, *P* < 0.05. *D*: Group averages of maximum PAD amplitudes (late peak for V3 PAD), * drPAD significantly larger, *P* < 0.05. *E*: Central latency of V3 PAD, drPAD and first spike evoked in V3 neurons (the latter as in Fig 4G, *n* = 26), * significantly different than V3 PAD latency, *P* < 0.05. *F-G*: Amplitude and peak time of the early and late peaks in V3 PAD, * late peak amplitude significantly larger than early peak, or V3 peak times significantly different than drPAD peak time, *P* < 0.05. *H*: Schematic of trisynaptic circuit underlying PAD with GABAergic inhibition of V3 neurons, explaining disinhibition of late PAD with gabazine. *I – J*: Changes in V3 PAD and drPAD with gabazine, * significant change in V3 PAD (*n* = 16) and drPAD (*n* = 21) amplitude and duration, *P* < 0.05.

Interestingly, the late peak of V3 PAD was largely blocked by gabazine (the portion which we refer to as V3 GABA PAD), as was most of the DR PAD, whereas the early peak of V3 PAD was unaffected, and so the former is mediated by GABA, and the latter is not. As mentioned, GABAA receptor antagonists like gabazine should disinhibit V3 neurons, and make their actions on PAD more pronounced, providing a duel excitatory and inhibitory action of gabazine (Fig 5H). Consistent with this, we found that as the peak of DR PAD or late peak of V3 PAD dropped in gabazine, there was a slow emergence of a very long lasting PAD with a duration much beyond the normal end of PAD (seconds; Fig 5B). Overall, in gabazine this left an early depolarization followed by a very long lasting PAD, both in V3 and DR evoked PAD. Thus, even the classic drPAD has a non-GABA mediated component that is revealed by disinhibiting the associated spinal circuits. Interestingly, even before adding gabazine, we see that the V3 PAD is a similar duration to the drPAD evoked by a strong DR stimulus adequate to activate cutaneous afferents (3xT), whereas the drPAD evoked by only lower threshold DR stimulation that mainly only activates proprioceptive afferents (2xT) is of shorter duration (Fig 5C), consistent with the V3 PAD perhaps involving a non-GABAegic PAD mediated by NMDA, since cutaneous afferent stimulation evokes a longer PAD (Lucas-Osma *et al*., 2018) that has recently been shown to be partly NMDA mediated (Zimmerman *et al*., 2019).

Considering that axonal NMDA receptors contribute to PAD in cutaneous afferents (Zimmerman *et al*., 2019) and that we find the action of V3 neurons may be faster than disynaptic, we next examined whether the glutamatergic V3 neurons themselves contact afferents and contribute to a glutamate-mediated PAD in Ia proprioceptive afferents. Indeed, we found that neurobiotin filled Ia afferents receive VGLUT2_+_ V3 innervation (Fig 6A). These V3 contacts were at branchpoints where nodes are located in large myelinated branch points of these afferents (at 51% of nodes examined) and at the terminals (at 36% of afferent terminal boutons examined), similar to the GABAergic innervation of these myelinated afferents that we have detailed previously (Hari *et al*., 2021).

**Fig 6.**
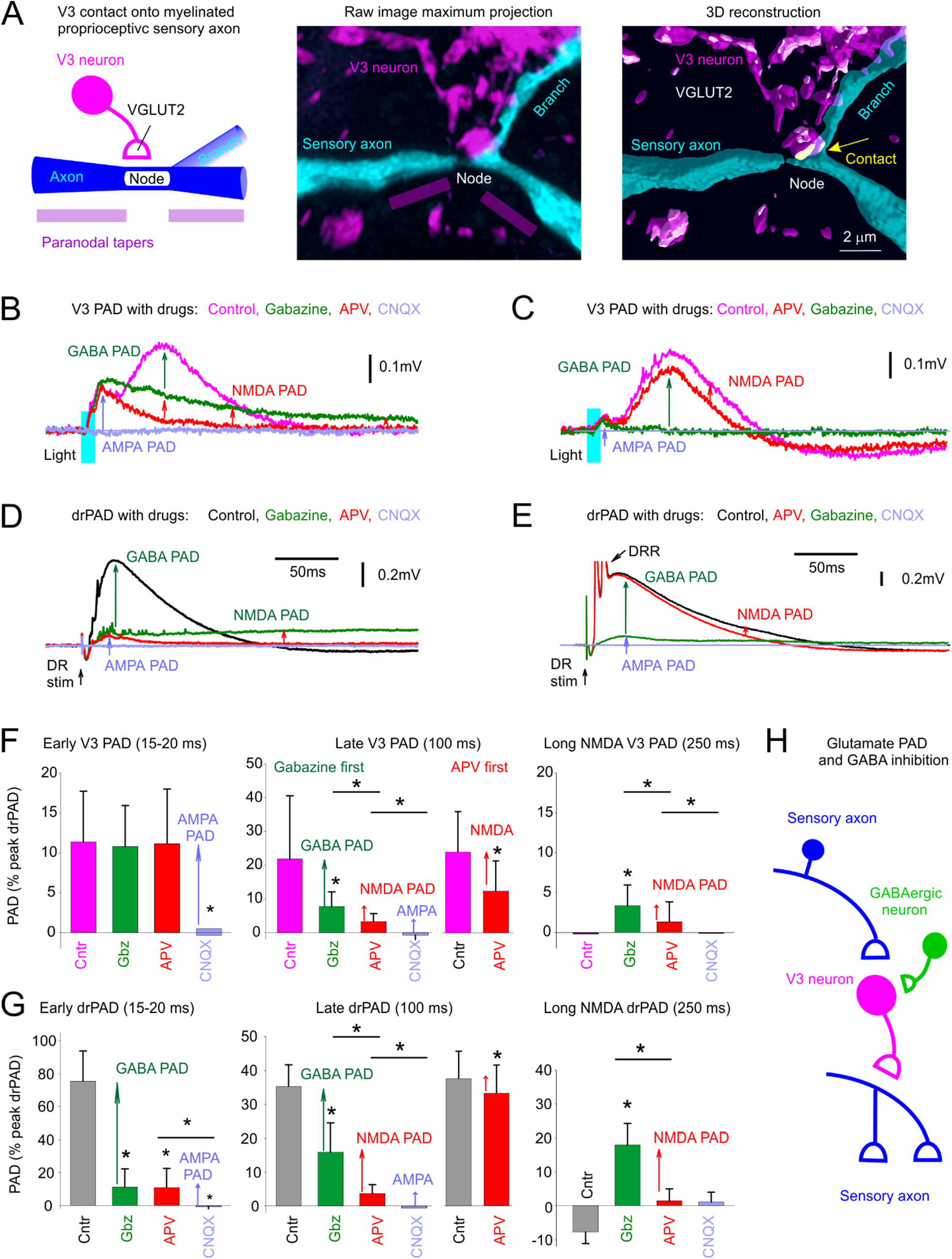
V3 neurons evoke a glutamate-mediated PAD, independent of GABA, by direct innervation of afferents. *A*: V3 neuron contact (VGLUT2_+_) on large myelinated Ia afferent branch in the intermediate spinal cord, at a branch point where the node of Ranvier is located (node also identified by the paranodal taper; afferent filled with neurobiotin). V3 neurons labelled with EYFP in Sim1//ChR2-EYFP mouse. Contact shown in yellow computed from 3D reconstruction on right. *B*: PAD evoked by a brief light pulse (10 ms pulse, λ = 447nm laser, 0.7 mW/mm_2_, 3xT, laser aligned as in Fig 4B) in Sim1//ChR2-EYFP mouse, and GABA PAD, NMDA PAD and AMPA PAD components estimated by the change with sequential application of gabazine, APV and CNQX, respectively. Recorded in S3 DR by grease gap (DRPs). *C*: Same as *B*, but APV applied first, with sequential application of APV, gabazine, and CNQX. *D-E*: Same as *B-C*, but for drPAD evoked by DR stimulation (Ca1 DR 0.1 ms, 2xT). *F*: Group averages of V3 PAD early peak (at 15-20 ms), late peak (at 100 ms) and long lasting NMDA PAD (at 250 ms), with sequential application of gabazine, APV CNQX (gabazine first condition) or APV first (only detailed for late V3 PAD since early PAD unaffected, not shown). * significant PAD reduction with drug application, P < 0.05, n = 20 gabazine first recordings, and n = 11 APV first recordings, from S4 and S3 DR combined. *G*: Group averages of drPAD amplitudes at same times as measured for V3 PAD in *F*, for comparison. * significant PAD reduction with drug application, *P* < 0.05, *n* = 20 gabazine first recordings, and n = 28 APV first recordings. *H*: Schematic of the putative disynaptic V3 neuron circuit mediating PAD independently of GABA, but with GABA tonically inhibiting V3 neurons, leading to a disinhibition of this glutamate-mediated PAD with gabazine.

Consistent with anatomical connections of V3 neurons to afferents, we found that the NMDA antagonist APV blocked the long-lasting component of both V3 PAD (Fig 6B, C, F) and drPAD (Fig 6D, E, G), indicating the presence of an APV-sensitive PAD mediated by NMDA (termed NMDA PAD). This NMDA PAD was best seen after applying GABA antagonists (gabazine first, Fig 6B, D, F, G), which again disinhibited the spinal cord circuits and so amplified the remaining PAD. However, APV also directly reduced PAD without gabazine present (Fig 6C,E-G), indicating some resting state NMDA PAD. Together these results support the existence of an NMDA-mediated PAD that is independent of GABA, and unmasked during the disinhibition of the spinal cord by gabazine. This NMDA PAD had a long onset latency (12.4 ± 7.1 ms) and lasted for up to 3 sec (Fig 5I-J).

After blocking GABA PAD with gabazine and NMDA PAD with APV there still remained an early rising short PAD (Fig 6B-G), corresponding to the early peak in the V3 PAD detailed above. This is likely also mediated by glutamatergic V3 neuron contacts on afferents, based on its very short latency (relative to the V3 spike latency), and is sensitivity to the AMPA/kainate receptor blocker CNQX, consistent with it being mediated by AMPA/kainate receptors (abbreviated AMPA PAD). Because CNQX also blocks afferent transmission, this PAD could alternatively be mediated by another transmitter like glycine or ACh, though antagonists to these receptors (strychnine or tubocurarine)(Shreckengost *et al*., 2021) did not reduce PAD (not shown). Thus, tentatively we call this AMPA PAD. AMPA PAD is seen in V3 PAD (Fig 6B,C), as well as drPAD (Fig 6D, E), but the latter AMPA drPAD is small (Fig 6D,E) and was likely previously missed in studies of drPAD. Taken together, AMPA PAD and NMDA PAD are likely mediated by a common glutamate innervation of afferents, in a minimally disynaptic circuit depicted in Fig 6H.

### Radiating intersegmental PAD caused by long propriospinal V3 axons

V3 neurons are propriospinal and commissural neurons with long axons that ascend and descend the spinal cord in the ventral and ventrolateral white matter (Fig 7A) (Zhang *et al*., 2008; Blacklaws *et al*., 2015). Thus, these long projecting axons may well help explain the characteristic radiating nature of PAD, where one nerve or dorsal root stimulation can activate PAD in many muscle afferents, including in afferents many segments away and across the midline (Lucas-Osma *et al*., 2018). In contrast, GABAergic GAD2_+_ neurons involved in PAD are small interneurons that cannot explain this radiating PAD (Rudomin & Schmidt, 1999; Betley *et al*., 2009). To examine the details of how V3 neurons activate PAD across spinal segments, we recorded PAD from a given dorsal root while selectively activating V3 neurons at varying spinal segments on one side of the cord, by focal light application in Sim1//ChR2 mice. Overall, the GABA-mediated late peak of V3 PAD was best evoked by light applied at or below, but not above, the DR where PAD was measured, and the peak was equally large ipsilateral and contralateral to the light (Fig 7B, D, E). This is consistent with GABA PAD being mediated by ascending V3 propriospinal tracts in the sacral cord, as depicted by a single ascending axon in the schematic of Fig 7A. Since ChR2 is expressed all along the propriospinal V3 axons as well as at the cell body (seen with EYFP expression in Sim1//ChR2-EYFP mice; not shown), the finding that light applied above a given root does not evoke a large V3 PAD in the root indicates that spikes evoked in ascending propriospinal axons by light do not travel antidromically back down to parent V3 neurons at or below the root, otherwise there should be a similar PAD with light above or below the root. This lack of antidromic spike conduction in ascending propriospinal axons is consistent with our observation that a rostral activation of these V3 axons produces a spike propagation failure in the antidromic direction, as detailed above (Fig 4E).

**Fig 7.**
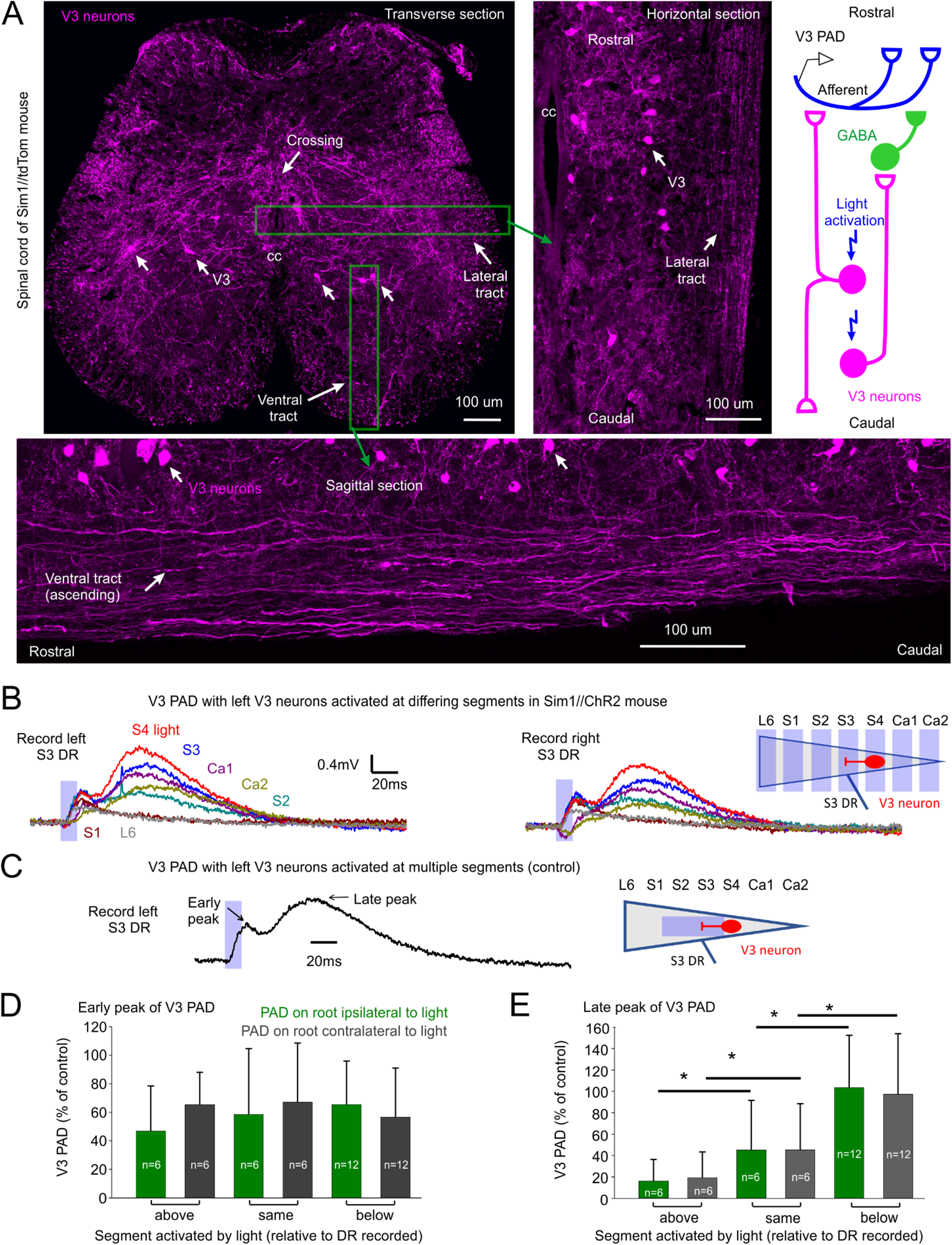
Intersegmental propriospinal projections mediating PAD. *A*: V3 neurons in sacral S3 spinal cord of Sim1//tdTom mice shown in transverse (top left, repeated from Fig 4A), sagittal (bottom) and horizontal (right) planes (the latter two at approximate locations indicated by green boxes), showing axonal tracts in the white matter formed by V3 neurons. Also, some V3 axons crossed the midline, as indicated. Schematic of proposed axonal projections of V3 neurons, with V3 neurons involved in GABA PAD (late peak of V3 PAD) mainly ascending from the lower sacral V3 neurons, and V3 neurons involved in glutamatergic PAD (AMPA PAD, early peak) both ascending and descending. *B-C*: V3 PAD in sacral S3 DRs evoked by light focused on the left lateral edge of the cord, at varying sacral and caudal segments indicated (*B*, by turning the narrow columnated beam of Fig 4B across the cord). V3 PAD was similar in left and right S3 DRs (commissural). Note that S3 PAD evoked in the S3 DR by applying light at S4 (red) was largest, sometimes even larger than PAD evoked by light applied across multiple segments at and above the S3 level (*C*; with columnated light turned to align with cord over S2-S3, as was usual arrangement elsewhere; Fig 4B), the latter used as our control PAD with which to normalized responses. *D-E*: Normalized group averages of early and late peaks of V3 PAD, grouped by whether the light was applied above (rostral to), at, or below (caudal to) the segment of the root where PAD was recorded (S3 or S4 DRs). * significant difference with applied light above or below root segmental level, P < 0.05, n = 6 – 12 roots per condition.

Interestingly, the glutamate-mediated early peak of V3 PAD in a given root was equally large when we activated V3 neurons with light above, at, or below the segment of the root (Fig 7B, D), suggesting that both ascending and descending V3 neurons cause this glutamate mediated PAD, and consistent with a differing underlying circuit, as depicted by a bipolar V3 neuron on Fig 7A schematic. In contrast, direct activation of GABAergic GAD2 neurons in GAD2//ChR2 mice led to a PAD that was mainly only large in roots directly under the light application site (not shown), consistent with the small special extent of these neurons (< 1 mm)(Hughes *et al*., 2005).

### V3 neurons facilitate proprioceptive sensory transmission to motoneurons

We next turned to examining the functional action of V3 neurons on sensory transmission. Recently, we have shown that PAD acts to directly facilitate spike transmission in afferents, including both antidromically and orthodromically propagated spikes (Lucas-Osma *et al*., 2018; Hari *et al*., 2021). Considering that our ArchT inhibition of V3 neurons demonstrated that at least half of drPAD is mediated through V3 action, we returned to this ArchT data to examine whether V3 neurons also contribute to spike propagation (Fig 3A). For this, we examined spikes produced in afferents by drPAD, which propagate both to the motoneurons and antidromically out the DR the latter which we measured (termed the dorsal root reflex, DRR, Fig 8A-B, 3A)(Lucas-Osma *et al*., 2018). These spikes were markedly reduced in incidence (Fig 8B) and on average reduced by half with the ArchT inhibition of V3 neurons (Fig 8C), consistent with a similar reduction in drPAD itself. Thus, V3 neurons facilitate spike transmission in sensory axons, likely via their depolarizing action (PAD) helping sodium channel spike initiation, similar to the general function of PAD (Hari *et al*., 2021).

**Fig 8.**
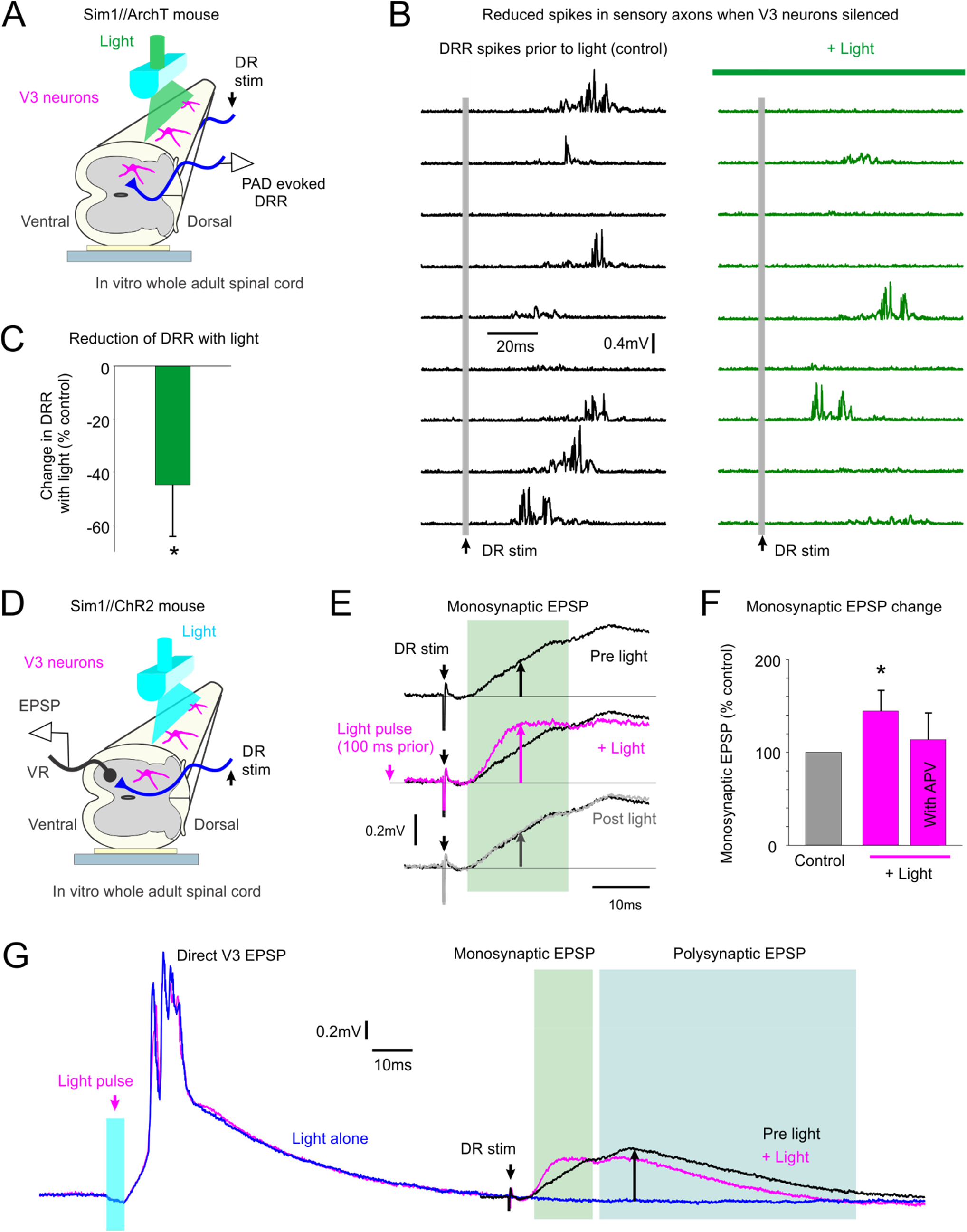
V3 neurons increase spike transmission in sensory afferents. *A-B*: Dorsal root reflex (DRR) recorded in S4 DR from stimulating the Ca1 DR (0.1 ms, 2xT; raster plot of 9 trials at 0.1 Hz) in Sim1//ArchT mice, before and after silencing V3 neurons with light (532nm, 5 mW/mm_2_). The drPAD evoked by this stimulation is shown in Fig 3A (same recording), where the rising phase of PAD produced the DRR, but here the DRR is shown in isolation by removing the slow PAD response with a 100 Hz high pass filter and then rectifying. *C*: Change in integrated area under rectified DRR with silencing of V3 neurons, * significant change, P < 0.05, n = 8. *D-E*: Motoneuron monosynaptic EPSP evoked by dorsal root stimulation (S4 DR, 0.1 ms pulse, 1.1xT) measured with grease gap recordings from ventral root (S4 VR, giving composite EPSP of motoneuron pool; top trace in *E*). Light activation of the V3 neurons (10 ms pulse, λ = 447nm laser, 0.7 mW/mm_2_, 3xT, laser aligned as in Fig 4B) applied 100 ms prior to the same DR stimulation increased the resulting monosynaptic EPSP (magenta trace in *E*). *F*: Group averages of change in monosynaptic EPSP with light, both without (as in *E*) and with APV (50 µM) in the bath. * significant increase, P < 0.05, n = 6 without APV, n = 4 with APV present, in *F*. *G*: Same data as in *E*, but on longer time base to show direct response to the V3 activation (left) and decrease in polysynaptic reflex evoked by the DR stimulation (right).

We also examined changes in orthodromic afferent spike transmission induced by V3 neurons, by examining the monosynaptic transmission of Ia afferents to motoneurons, with direct recordings from motoneurons while evoking monosynaptic EPSPs with DR stimulation pulses (Fig 8D-E). When we activated V3 neurons optogenetically in Sim1//ChR2 mice to produce a V3 PAD, the monosynaptic EPSPs in motoneurons was as expected increased during the V3 PAD (Fig 8E-F). Thus, the V3 PAD increases proprioceptive afferent transmission to motoneurons. This facilitation was somewhat reduced when APV was applied to block the NMDA receptors and associated NMDA PAD, though further experiments are needed to confirm this (with higher n).

These monosynaptic EPSP experiments were somewhat confounded by the strong postsynaptic EPSPs that V3 neurons themselves directly produced in motoneurons, which lasted about 50 - 100 ms (Fig 8G)(Lin *et al*., 2019). Thus, to avoid postsynaptic actions of these long motoneuron EPSPs, we tested the DR-evoked monosynaptic EPSP at 100 ms after the light application, a time when V3 PAD was still present (Fig 5C, PAD lasts ∼150 ms), but the V3 evoked motoneuron EPSP had subsided. Thus, the facilitation of the monosynaptic EPSP by V3 neurons at this time (Fig 8E-F) is likely from the V3 neuron actions on the sensory axons, and not its postsynaptic action on motoneurons. Nevertheless, the polysynaptic pathways directly activated by V3 neurons likely became refractory to subsequent reactivation, since we observed that DR-evoked polysynaptic ESPS decreased with light (Fig 8G), though we did not quantify this more complex process. To complicate matters further, possibly V3-evoked PAD and DRRs contribute to the polysynaptic EPSPs evoked by light, with the latter leading to post activation depression of the monosynaptic afferent transmission (Hari *et al*., 2021), though this remains to be explored.

### Increased glutamate PAD and decreased GABA PAD after SCI

Considering that glutamate dependent PAD is unmasked by bringing up the excitability of spinal circuits with gabazine, we wondered whether the marked increase in spinal cord excitability with chronic a spinal cord injury (SCI, sacral S2 transection)(Murray *et al*., 2010; Lin *et al*., 2019) also unmasked glutamate-mediated PAD. For this we examined Sim1//ChR2 mice with an S2 sacral transection ∼1 – 1.5 months previously, and measured NMDA, AMPA and GABA PAD components with drug applications (APV, gabazine and CNQX-sensitive components, respectively; Fig 9A-D). Interestingly, the absolute size of the drPAD and main peak of the V3 PAD (late peak) did not change with SCI, whereas the small glutamate-dependent early peak of the V3 PAD markedly increased with SCI (Fig 9E, F). Further, the balance of GABA and glutamate PAD components shifted markedly to increased glutamate and less GABA contribution with SCI, as seen by normalizing these components as a fraction of the peak PAD recorded for each root (pre-drug drPAD; Fig 9G). For the V3 PAD evoked by light, its APV-sensitive NMDA PAD component made up a larger proportion of the total PAD after SCI, at times larger than GABA PAD (Fig 9 A-C, G). This occurred regardless of whether (Fig 9B) or not (Fig 9D) we added gabazine prior to APV, indicating that there is a large NMDA PAD in the resting cord with SCI (without gabazine), unlike without injury, suggesting increased direct V3 action on afferents. In contrast, NMDA PAD made up only a minor faction of V3 PAD without injury, even when measured after disinhibition with gabazine (Fig 9A, G). AMPA PAD likewise made up a larger fraction of the total PAD after SCI (Fig 9A-D, G; measured after NMDA and GABA receptor blockade), and had a longer duration. In contrast, the gabazine-sensitive GABA V3 PAD decreased with SCI (Fig 9G). However, this decrease only reached significance when APV was given prior to gabazine, rather than the reverse, because gabazine alone increased the NMDA PAD at the same time it blocked the GABA PAD, decreasing the overall gabazine-sensitive measurement.

**Fig 9.**
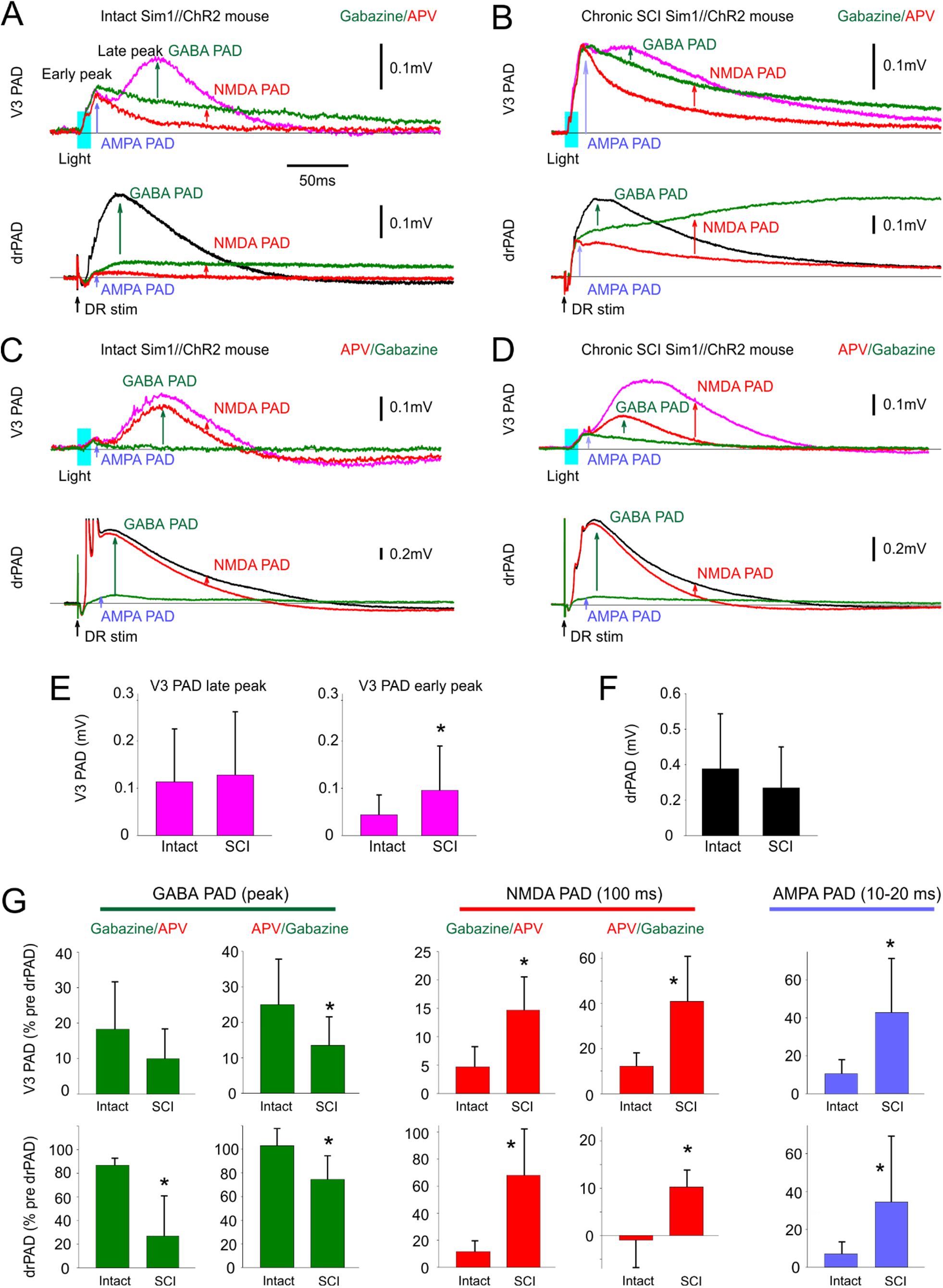
Chronic spinal cord injury (SCI) increases NMDA and AMPA PAD while reducing GABA PAD. *A*: Changes in V3 PAD and drPAD recorded in S3 DR (as detailed in Fig 6B and D) with sequential application of gabazine and APV in Sim1//ChR2 mouse. GABA and NMDA PAD measured by drug induced reductions, and AMPA PAD measured as remaining PAD in both drugs. V3 PAD and drPAD evoked by a brief light or DR stimulation, respectively (as detailed in Fig 6B and D). *B*: Same as *A* but in Sim1//ChR2 mouse with an S2 spinal transection 1 month previously (chronic SCI). *C-D*: Same as *A-B*, but with APV added before gabazine to estimated resting state NMDA PAD. *E-F*: Group averages of changes in PAD with SCI, with significant change (*) in early peak of V3 PAD, but not the late peak of V3 PAD or drPAD, *P* < 0.05, n = 31 and 15 from control and SCI mice. *G*: Group averages of changes in GABA PAD (at peak), NMDA PAD (recorded at 100 ms latency) and AMPA PAD (recorded at peak between 10 – 20 ms) with SCI, * significant change, P < 0.05, n = 20 and 6 with gabazine given first, without and with SCI respectively, and n = 12 and 6 with APV given first, without and with SCI respectively. All changes with drugs normalized by predrug drPAD peak.

The GABA and glutamate components of the drPAD showed similar changes to the V3 PAD (Fig 9A-D). In this case GABA drPAD made up >85% of the total peak drPAD without injury, whereas after SCI this decreased to either 35 or 75%, depending on whether or not APV was present prior to adding gabazine to measure the GABA PAD (Fig 9G), where again APV decreased cord excitability, leading to less PAD. In contrast, the NMDA and AMPA drPAD components made up a larger longer lasting proportion of the total drPAD after SCI (Fig 9G). Even the small or absent NMDA PAD measured without disinhibiting cord with gabazine increased with SCI (Fig 9G), again consistent with SCI leading to an increased excitability of the V3 circuits that directly produce PAD.

## DISCUSSION

Our results demonstrate that propriospinal V3 neurons play a major role in depolarizing group Ia proprioceptive sensory afferents across many spinal segments, facilitating afferent spike propagation, and ultimately modulating proprioceptive sensory transmission to motoneurons. They do this by both GABA-dependent and GABA-independent pathways. Specifically, V3 neurons appear to serve as the first order neurons in the classic trisynaptic circuit that causes GABA-mediated PAD in Ia afferents in response to proprioceptive sensory stimulation (Jankowska *et al*., 1981; Rudomin & Schmidt, 1999), since silencing V3 neurons eliminates the majority of this PAD (drPAD), and these neurons have the appropriate anatomical and functional connections. That is, V3 neurons receive extensive sensory innervation, including proprioceptive Ia afferent input, project over many segments, and in turn activate GAD2_+_ GABAergic neurons that are known to form axoaxonic connections back onto Ia sensory afferents. This circuit modulates sensory transmission to motoneurons, as evidenced by the action of PAD evoked by optogenetic activation of V3 neurons (V3 PAD), and importantly acts over many segments, explaining the radiating nature of PAD and its reflex modulation (Barron & Matthews, 1938; Eccles *et al*., 1961; Eccles *et al*., 1962; Rudomin & Schmidt, 1999; Lucas-Osma *et al*., 2018). Whether part of the V3 PAD we measured occurs in low threshold afferents (LTMRs, i.e., cutaneous afferents), in addition to Ia afferents, is unknown as we did not specifically examine these afferents.

Glutamatergic CCK+ interneurons serve as first order neurons in the GABA-dependent PAD evoked in LTMRs (Zimmerman *et al*., 2019), but unlike V3 neurons, are not propriospinal neurons, having only limited extent of synaptic contacts confined to the LTMR recipient zone (Abraira *et al*., 2017). Thus, it remains to be determined whether the radiating intersegmental nature of sensory evoked PAD observed in cutaneous afferents (Eccles & Krnjevic, 1959) is also mediated by a propriospinal neuron like sensory-evoked PAD in proprioceptive afferents.

Furthermore, we unexpectedly found that V3 neurons cause a pronounced GABA-independent PAD mediated by NMDA and AMPA/kainate receptors (NMDA and AMPA PAD), at least partly mediated by direct connections from V3 neurons to Ia afferents. This enables sensory stimulation to disynaptically activate a PAD, faster than the classic trisynaptic circuit that mediates GABA PAD. NMDA receptors and associated RNA are known to be expressed in low and high threshold mechanoreceptors and proprioceptors, including Ia afferents (Wu *et al*., 2021)(Dedek, 2022).

Presynaptic NMDA receptors (preNMDA) produce PAD in LTMR afferents (Zimmerman *et al*., 2019), and these preNMDA receptors have been shown in other systems to facilitate synaptic transmission (Dedek, 2022). This NMDA PAD in LTMRs is mediated by small dorsally oriented glutamatergic VGLUT3_+_ interneurons, and thus are again different from the large propriospinal V3 neurons that we observe mediate NMDA PAD (Peirs *et al*., 2015; Zimmerman *et al*., 2019). Whether a similar preNMDA arrangement exists in proprioceptive afferents is unknown since we did not examine NMDA receptors distribution on Ia afferents, though at least we know that these afferents are innervated by V3 neurons that cause a direct fast glutamate-dependent PAD, and thus could contribute to the increased sensory transmission we observed with V3 activation (which was somewhat APV sensitive). Alternatively, NMDA receptors on neurons in polysynaptic pathway could also contribute to the NMDA PAD we observed. Indeed, likely, all forms of PAD (GABA, NMDA, and AMPA PAD) are likely additionally mediated by longer latency polysynaptic pathways than the minimally trisynaptic and disynaptic pathways that we describe here. For example, the complex central pattern generator (CPG) circuits that underlie walking and the corticospinal tracts involved in grasping are known to evoke PAD (Anden *et al*., 1966; Rossignol *et al*., 2006; Ueno *et al*., 2018; Lalonde & Bui, 2020; Moreno-Lopez *et al*., 2021), and thus long-loop circuits may evoke the differing forms of PAD via V3 neurons, and could partly explain the long lasting NMDA PAD (via the CPG). Furthermore, we cannot rule out fast AMPA PAD triggering spikes in afferents (DRRs) that in turn trigger further GABA PAD reverberatively in other afferents, and perhaps explaining the delay in the peak of the GABA component in the V3 PAD.

Under resting in vitro conditions NMDA and AMPA drPAD evoked by sensory stimulation is not very large, at least not the many second long component of NMDA PAD, explaining why it has not been well studied in the past. However, direct activation of V3 neurons evoke more pronounced NMDA and AMP PAD, suggesting that any system, including the CPG, that engages the V3 neurons (Zhang *et al*., 2008) may also evoke pronounced GABA-independent NMDA and AMPA PAD. Indeed, with increased spinal cord excitability, caused by either blocking GABA receptors or chronic spinal cord injury, both NMDA and AMPA PAD become much more important. Whether or not such glutamate-mediated PAD is important in awake animals remains an open question, though it may be best activated via cutaneous stimulation, since it seems to be associated with cutaneous evoked PAD (with stronger stimulation), and similar NMDA-associated PAD has been reported in other in vitro systems in their resting state (Russo *et al*., 2000; Lucas-Osma *et al*., 2018; Zimmerman *et al*., 2019). Furthermore, it may well be that during complex movements like locomotion, where PAD is increased by the CPG, NMDA PAD may play a larger role.

The duel action of GABA on PAD that we uncovered is particularly relevant to understanding studies of PAD that employ GABA blockers. That is, GABAA receptors on the V3 neuron inhibit these neurons and their PAD action, whereas the GABAA receptors on afferents are indirectly activated by V3 neuron connection to GABAergic neurons. Thus, in our and other previous studies (Fink, 2013; Lucas-Osma *et al*., 2018; Hari *et al*., 2021) where GABAA receptors blockers are used to quantify GABA’s role in PAD and sensory transmission, there is both a direct block of GABA PAD and a disinhibition of V3 neurons, the later which enhances NMDA and AMPA PAD. Thus, the size of the PAD mediated by afferent GABAA receptors or its underlying functional action may be underestimated. Perhaps a better estimate of GABA PAD is obtained by first blocking NMDA receptors (and NMDA PAD), prior to blocking GABAA receptors, as we previously also noted when evaluating the actions of GABAA antagonists on monosynaptic transmission (Hari *et al*., 2021).

While GABA-mediated PAD (GABA PAD) is a major focus of this paper, GABA PAD can be viewed as a proxy measure for more general GABAergic action on afferents, regardless of whether that action produces PAD. That is, axoaxonic innervation of proprioceptive afferents by GABAergic neurons has a two opposing actions: causing presynaptic inhibition of sensory transmission by activating GABAB receptor at afferent terminals that do not produce PAD, and causing a facilitation of sensory action conduction by activating GABAA receptors at or near nodes of Ranvier that cause PAD, which in turn assists local sodium spike conduction at the nodes (called nodal facilitations) (Hari *et al*., 2021). Thus, our finding that V3 neurons mediate GABA PAD suggests that they may be involved in both presynaptic inhibition and well as nodal facilitation. So far, we have only shown that V3 neuron action facilitates monosynaptic sensory transmission to motoneurons, consistent with them at least causing nodal facilitation. Whether they also produce GABAB mediated actions, including the well known rate dependent depression (RDD) of the monosynaptic reflex (Lev-Tov *et al*., 1988; Hultborn *et al*., 1996), remains an open question.

Our demonstration that V3 neurons still cause PAD after chronic spinal transection suggests that descending innervation from the higher spinal or supraspinal tracts are not needed for V3 production of PAD. Indeed, we found that at least in the sacral cord it was ascending intersegmental propriospinal V3 neurons that best evoked GABA PAD. Interestingly, the early glutamate-dependent PAD was evoked equally well by ascending or descending intersegmental V3 neurons, suggesting differing underlying population mediating GABA PAD compared to AMP PAD, though this remains to be examined anatomically (Blacklaws *et al*., 2015). With spinal cord injury the AMPA and NMDA PAD increased markedly, whereas the GABA PAD decreased, again suggesting different circuits underling these types of PAD. Interestingly, the overall PAD did not change much with injury, just the balance of GABA and glutamate dependent PAD, consistent with previous observations of not much change in PAD with SCI, except for a higher sensory threshold to be evoked (Caron *et al*., 2020). The latter may be explained by the increased NMDA PAD, which seems to be evoked by higher threshold cutaneous afferents, compared to low threshold Ia afferents, as detailed above. The functional outcomes of the increased NMDA PAD with SCI remains to the determined, though it may be a compensation for loss of spinal GABAergic neurons with SCI (Meisner *et al*., 2010).

Importantly, the long propriospinal projections of V3 neurons (Blacklaws *et al*., 2015) allows local sensory or V3 neurons stimulation to evoke PAD in afferents in distant segments and across the midline, thus explaining the remarkable radiating nature of PAD that has been a mystery for nearly a century (Barron & Matthews, 1938; Eccles & Krnjevic, 1959). This is on contrast to the small GABAergic neurons involved in PAD that only locally affect PAD (Rudomin & Schmidt, 1999; Betley *et al*., 2009).

Further, the intrinsic properties of V3 neurons, including Na PICs, allow them to respond with very long plateau-like depolarizations following a brief activation, thus explaining the long-lasting nature of PAD, and its potent action on modulating sensory transmission over long time periods. Interestingly the somewhat TTX resistant nature of the sodium channels in V3 neurons, suggests that that the previous observation that an NMDA PAD triggered by high threshold afferents (C fibre) is TTX resistant is not just explained by TTX resistant channels in C fibres themselves (Russo *et al*., 2000; Lucas-Osma *et al*., 2018). Instead, it is likely also explained by TTX resistant sodium channels in V3 neurons and the direct connects of V3 neurons to afferents that mediates NMDA PAD. This implies that the previously supposed microcircuit underlying this C fibre mediated PAD may not actually be a microcircuit, but instead a large scale circuit where PAD is evoked by C fibre inputs onto V3 neurons that in turn monosynaptically dopolarize large myelinated sensory afferents, like Ia afferents. This is consistent with the intersegmental nature of this TTX resistant PAD, occurring in one dorsal root in response to stimulation of another, and likely mediated by the propriospinal axons of V3 neurons (Russo *et al*., 2000; Lucas-Osma *et al*., 2018). However, the actual details of C fibres innervation of V3 neurons and the TTX resistant sodium channels in V3 neurons remains to be explored to confirm these ideas.

Taken together these properties of V3 neurons allows them to form wide ranging spatial temporal integration of their sensory and other inputs (CPG) to control PAD and sensory transmission.

V3 neurons have previously been studied in the context of locomotion and general motoneuron function (Zhang *et al*., 2008), and so our finding that they play a major role in sensory transmission needs to be reconciled with their motor functions. The underlying commonality may be that V3 neurons act like neuromodulators with their slow integrator properties, turning on and off sensory and motor systems during differing tasks. For example, we previously found that V3 neurons cause direct sustained depolarizations of motonenurons, and seem to be crucial in turning on sustained locomotor-like activity and spasms, especially after spinal cord injury in mice (Lin *et al*., 2019). Further, in zebra fish V3-like neurons steadily depolarize during bouts of locomotion, and are not rhythmically active (Bohm *et al*., 2022). Instead, they serve to amplifying and prolonging motor output, consistent with them causing a broad depolarization of the many CPG neurons and motoneurons they innervate, and thus acting as high-level neuromodulators that amplify and integrate of activity, rather than controlling fine rhythm details (Bohm *et al*., 2022). More generally, the long ascending and descending propriospinal tracts of V3 neurons have been suggested to be involved in coordinating reciprocal flexor-extensor and left-right multi-joint muscle activity during locomotion, though they somehow do this without controlling the details of the movements (Zhang *et al*., 2008; Bohm *et al*., 2022). That is, V3 neurons and more generally other propriospinal neurons, are not normally needed for basic locomotor rhythm generation (i.e., silencing V3s does not stop the CPG or walking) and are often not rhythmically driven by the CPG (Zholudeva *et al*., 2021). Instead, they may be involved in the higher level task of choosing which general motor action occurs, whether it is reciprocal stepping, hopping or swimming (Zhang *et al*., 2008; Zholudeva *et al*., 2021). With this view that V3 neurons act like neuromodulators that ready the motor system for movement, its makes sense that V3 neurons also modulate sensory neurons, readying sensory feedback during movement, similar to how the Jendrasik maneuver enhances sensory feedback via the monosynaptic stretch reflex (Zehr & Stein, 1999). Whether subpopulations of V3 neurons target these differing sensorimotor actions remains an open question (Borowska *et al*., 2013; Chopek *et al*., 2018; Deska-Gauthier *et al*., 2018), though this would allow flexibility in controlling sensory function during movement. In retrospect, the view that V3 neurons broadly supervise sensorimotor systems, taken together with V3 neurons causing PAD, suggests that PAD is just one of the many actions of V3 neurons involved in coordinating complex movements, and PAD is not part of an isolated system regulating sensory transmission.

